# AdaLiftOver: High-resolution identification of orthologous regulatory elements with adaptive liftOver

**DOI:** 10.1101/2022.06.03.494721

**Authors:** Chenyang Dong, Sündüz Keleş

**Affiliations:** Department of Statistics; Department of Biostatistics and Medical Informatics

**Keywords:** Orthologous mapping, Genome-wide association studies, Model organism, Comparative epigenomics, Transfer learning, LiftOver

## Abstract

Elucidating orthologous regulatory regions for human and model organism genomes is critical for exploiting model organism research and advancing our understanding of results from the genome-wide association studies. Sequence conservation is the *de facto* approach for finding orthologous non-coding regions between human and model organism genomes. However, existing methods for mapping non-coding genomic regions across species are challenged by the multi-mapping, low precision, and low mapping rate issues. We develop Adaptive liftOver (AdaLiftOver), a large-scale computational tool for identifying orthologous non-coding regions across species. AdaLiftOver builds on the UCSC liftOver framework to extend the query regions and prioritizes the resulting candidate target regions based on the conservation of the epigenomic and the sequence grammar features. Evaluations of AdaLiftOver with multiple case studies, spanning both genomic intervals from epigenome datasets and GWAS SNPs yield AdaLiftOver as a versatile method for deriving hard-to-obtain human epigenome datasets as well as reliably identifying orthologous loci for GWAS SNPs. The R package AdaLiftOver is available from https://github.com/ThomasDCY/AdaLiftOver.

## 1 Introduction

Genome-wide association studies (GWAS) have revealed many non-coding SNPs for complex human traits (Welter et al., 2014; Gallagher and Chen-Plotkin, 2018). However, identifying the effector genes of non-coding SNPs and elucidating their specific roles in disease etiologies are key challenges of modern GWAS. Model organism studies are important and underexploited resources for dissecting GWAS SNPs by experimentally perturbing the orthologous model organism loci for the human genomic regions of interest. Reliable maps of non-coding genomic regions between human and model organism genomes will not only improve our understanding of the evolution of regulatory mechanisms but also pinpoint orthologous regulatory elements for comparative genomics and epigenomics analysis.

Sequence alignment has made fundamental contributions to phylogenetic analysis and evolutionary biology (Earl *et al*., 2014). Leveraging DNA sequences as the mapping units is the standard approach to establish putative orthologous mappings across different species. The current architecture of translating genomic coordinates across genome assemblies is largely based on UCSC’s chained and netted sequence alignment results, which are summarized as chain files. The UCSC liftOver tool (Hinrichs *et al*., 2006) is the *de facto* mapping strategy in cross-species studies. More recently, bnMapper (Denas *et al*., 2015), which is a Python implementation similar to UCSC liftOver but leverages reciprocal chain files allowing for only one-to-one mappings across species has emerged. However, there are a number of practical drawbacks of these strictly sequence alignment-based mapping approaches of non-coding sequences. We group these into three categories as follows using the mappings between promoters of orthogolous human and mouse genes:

1. *The prevailing multi-mapping issues.* A given non-coding region in the human genome can be mapped, i.e., lifted over, to multiple mouse regions. For example, when we consider the 16,374 human genes with mouse orthologues, 94.7% of their promoters map to multiple mouse regions with an average of 38.8±22.0 regions (Supplementary Materials Section 1.1). Merging of the small gaps less than 10 bp yields mapping of the human promoters to an average of 3.68±2.73 mouse regions and still leaves 78.6% as mapping to multiple regions. In particular, 34.8% of the human promoters map to multiple mouse regions separated apart by at least 200 bp.
2. *Inaccurate mappings and low precision issues.* Sequence-based mapping of the 16,374 orthologous human and mouse promoters is prone to generating 2,849 (17.4%) false positive and false negative cases (Supplementary Materials Section 1.1). Fig. 1a illustrates an example of true positive mapping (76.3% of all orthologous promoters), where orthologous chain segments map the promoter region of *FEZF2* in human to that of *Fezf2* in mouse. In contrast, Fig. 1b displays a potential discrepancy, where the mouse counterpart of orthologous chain segments reside at the promoter region of *Opn4* while the human counterpart is located 2 kb away from the transcription starting site of *OPN4*. Consequently, both the UCSC liftOver and the bnMapper fail to map the *OPN4* promoter to mouse genome and create a potential false negative. We note that the false negative might become a false positive if we map from *Opn4* promoter in mouse to the human genome.
3. *Low mapping rates.* Unlike the highly conserved orthologous promoters, Cheng *et al*., 2014 found that ~50% of the transcription factor occupied regions failed to map to the mouse genome and Dong et al., 2021 observed that ~80% of diabetes related human GWAS SNPs were unmappable to the mouse genome. This highlights the general challenge of mapping human non-coding regions to model organism genomes.

**Figure 1:**
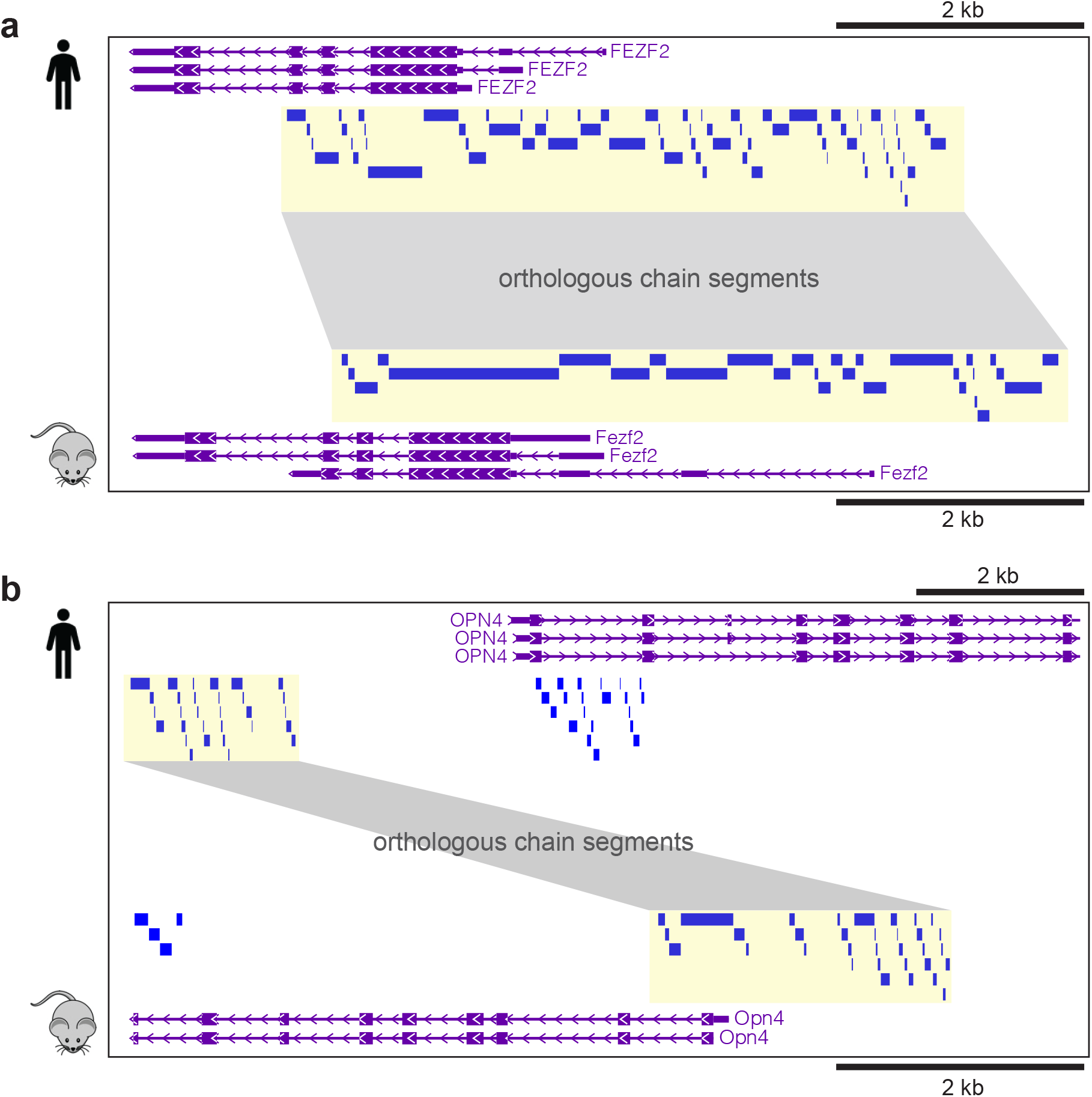
**a.** The promoter of *FEZF2* in human maps directly to the promoter of *Fezf2* in mouse. **b.** The promoter of *OPN4* in human maps indirectly to the promoter *Opn4* in mouse by allowing for a local window. The blue rectangles depict chain segments from the pairwise sequence alignment. Orthologous chain segments between human and mouse are highlighted in translucent yellow and connected with gray bands.

Recent deep learning applications have yielded advanced investigations of the regulatory code of DNA sequences. Early applications of deep convolutional neural network models used only DNA sequences to predict protein binding, histone modification, and chromatin accessibility (Zhou and Troyanskaya, 2015; Alipanahi *et al*., 2015). Basset (Kelley *et al*., 2016) predicted the impact of noncoding variants on cell type specific DNase-seq profiles. With larger scale and finer resolution, Basenji (Kelley *et al*., 2018) incorporated distal interactions and predicted a much larger collection of epigenome profiles. ExPecto (Zhou *et al*., 2018) evaluated the tissue specific gene expression changes for mutations. Cross-species investigations with deep neural networks (Kelley, 2020; Minnoye *et al*., 2020) implicated a higher level regulatory code beyond strict sequence conservation as playing a significant role for predicting orthologous functional regions. Minnoye *et al*., 2020 discovered examples of orthologous enhancers that sequence-based methods failed to identify. Beyond sequence conservation, functional genomic annotations are important complementary information to determine orthology (Kwon and Ernst, 2021). Many studies have revealed the evolutionary landscape of genomes and epigenomes by comparing matched datasets across species (Villar *et al.*, 2015; Gjoneska *et al*., 2015; Vierstra et al., 2014; Cheng *et al*., 2014; Brawand *et al*., 2011; Odom *et al*., 2007). EpiAlignment (Lu *et al*., 2019) is the first method that incorporates both matched ChIP-seq experiments and DNA sequences as the mapping units to identify orthologues between human and mouse. However, EpiAlignment allows for binary encoding of only one matched pair of functional genomic datasets which provides limited information for discriminating a conserved epigenome against a random one.

In order to address the limitations of strictly sequence-based mapping of orthologous regions and leverage higher-order regulatory grammar embedded in DNA sequences, we developed Adaptive liftOver (AdaLiftOver). AdaLiftOver is built on the UCSC liftOver framework for identifying and prioritizing orthologous regions by leveraging functional epigenome information. It enables mapping genomic coordinates between any two species with chain files and at least one pair of orthologous epigenome datasets. AdaLiftOver takes as input query genomic regions, the UCSC chain file, and one or more orthologous epigenome datasets (Fig. 2). We curated a list of orthologues epigenome datasets between human and mouse for general use from the ENCODE resources (Moore *et al*., 2020). AdaLiftOver allows the users to adaptively incorporate additional matched datasets and adjust the contribution of these datasets to the mapping. For each query region, AdaLiftOver generates a curated list of candidate target re-gions and priorizes them with a score from a logistic model. The users can retrieve the final set of mapping regions by retaining only the top candidate target regions exceeding a score threshold (Fig. 2).

**Figure 2:**
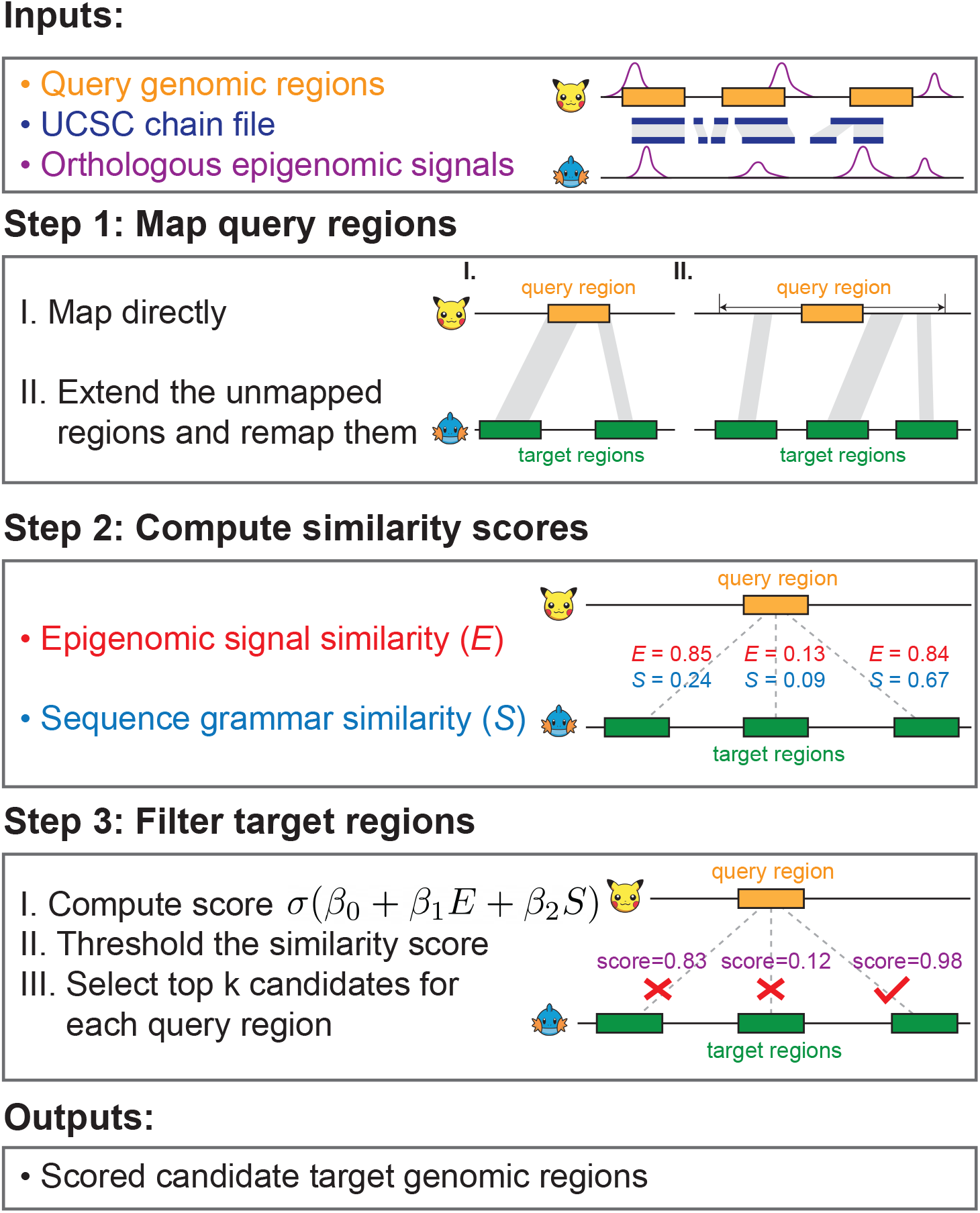
The AdaLiftOver workflow. The cartoon icons denote any two species with chain files. Top: query regions (orange) in the query genome; Bottom: target regions (green) in the target genome. Inputs: genomic coordinates of the query regions (orange), the UCSC chain file from the query genome to the target genome (blue), and the matched epigenome datasets (purple). Step 1: AdaLiftOver defaults to the UCSC liftOver if the query regions map successfully (I). If a query region does not map, AdaLiftOver extends the query region in a local window and applies the UCSC liftOver to this extended query region (II). AdaLiftOver merges small gaps among the resulting orthologous regions and generates candidate target regions based on these merged orthologous regions (indicated by translucent gray) with the same width as the query region. Step 2: AdaLiftOver uses local binary epigenomic and sequence grammar feature vectors to compute the similarity scores between the query region and each of the corresponding candidate target regions. Step 3: AdaLiftOver scores the candidate target regions with a logistic model (*β*_0_ = −3, *β*_1_ = 4, *β*_2_ = 5) based on their two similarity scores. The users can threshold these scores and rank the candidate target regions based on their probabilities of mapping to the query region. With score threshold of 0.4 or *k* = 1, AdaLiftOver picks the rightmost candidate target region with estimated probability of mapping as 0.98. Outputs: For each query region, AdaLiftOver outputs a scored and filtered list of candidate target regions that are most similar to the query region in terms of regulatory information.

We applied AdaLiftOver to a variety of case studies including genomic intervals (peaks) from ATAC-seq and ChIP-seq, and SNPs from GWAS datasets as queries. AdaLiftOver yields consistently superior performances than competing methods for mapping of both the peaks and the SNPs to the mouse genome. The R implementation for AdaLiftOver is available at https://github.com/ThomasDCY/AdaLiftOver.

## 2 Methods

### AdaLiftOver framework

AdaLiftOver is a large-scale computational framework that leverages functional regulatory information to enhance the UCSC liftOver. Specifically, AdaLiftOver implements a two-step strategy for mapping each query genomic region *Q* (Fig. 2), which could constitute genomic intervals from biochemically active regions of the genome (i.e., ChIP-seq peaks) or GWAS SNPs. We first directly apply the UCSC liftOver to map *Q.* If *Q* fails to map directly, we extend *Q* with a local window on both sides and then apply the UCSC liftOver on the extended query region to generate candidate target regions. Let *O*_1_ *O*_2_, ⋯, *O_m_* denote the resulting candidate regions. We note that it is possible to have no orthologous regions, i.e., *m* = 0. For each orthologous region *O_j_, j* = 1, ⋯, *m*, AdaLiftOver generates a list of evenly-spaced candidate target regions with a predefined resolution *T*_*j*,1_,*T*_*j*,2_, ⋯, *T_j,n_j__*, where *n_j_* ≥ 1 and their widths are set to be equal to that of *Q*. For simplicity, we denote all curated target genomic regions of *Q* as *T*_1_, *T*_2_, ⋯, *T_n_*, where 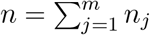. AdaLiftOver computes the local epigenomic feature vectors for *Q* and *T*_1_, *T*_2_, ⋯, *T_n_* as *e*_*Q*_ and *e*_*T*_1__, *e*_*T*_2__, ⋯, *e_T_n__*. Likewise, the local sequence grammar feature vectors are defined as *s_Q_* and, *s*_*T*_1__, *s*_*T*_2__, ⋯, *s_T_n__*. Then, the epigenomic and the sequence grammar feature similarities can be defined as *E_i_*, = sim(*e_Q_,e_T_i__*) and *S_i_*, = sim(*s_Q_,s_T_i__*), respectively, where *i* = 1,2, ⋯, *n* and sim(·) is a similarity function. AdaLiftOver scores each candidate target region *T*, with a logistic function *σ*(*β*_0_ + *β*_1_*E*_*i*_ + *β*_2_*S_i_*), where 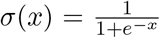 is the sigmoid function and (*β*_0_,*β*_1_,*β*_2_) are predefined logistic regression parameters estimable by the training module.

### Regulatory information similarity

#### Epigenomic features

We interrogated 67 matched ENCODE ChIP-seq and DNase-seq datasets between human and mouse from the following 10 tissues: heart, kidney, liver, lung, placenta, small intestine, spleen, stomach, testis, and thymus (Supplementary Materials Section 1.2). These datasets are integrated into AdaLiftOver and the users can augment these with additional epigenome datasets from matching tissues and/or orthologues cell types. While there are a number of ways to summarize the signal from epigenome datasets, due to the computational challenges we articulated in Supplementary Materials Section 1.4, we considered the local epigenomic features as 67-dimensional binary vectors from the overlap of the genomic regions with the peaks from the epigenome datasets. The choice of binarization provides a balance between the signal-to-noise ratio and the computational time, space, and memory requirements (Supplementary Materials Section 1.4). We also remark that, in all the case studies that follow, the query samples are not from these 10 tissues used to derive the epigenome features to illustrate robustness of AdaLiftOver for mapping query regions of interest without directly relevant epigenome datasets.

#### Sequence grammar features

We utilized 841 core vertebrate JASPAR motifs (Castro-Mondragon *et al*., 2022) as a list of “words” capturing the high-level “grammar” encoded by DNA sequences and are beyond traditional sequence alignment. We used “motifmatchr” (Schep *et al*., 2017) for fast motif scanning in the vicinity of genomic regions instead of storing and querying the genome-wide motif occurrences. We define the sequence grammar feature of a query region as the 841-dimensional binary vector.

#### Similarity metrics

For a pair of binary vectors *u,v* ∈ ℝ^*d*^, their weighted cosine similarity with weight *w* ∈ ℝ^*d*^ can be computed as:

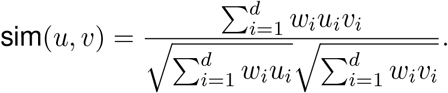

As default, AdaLiftOver uses equal weights while computing similarity scores. The users can specify the weights for computing the epigenomic feature similarities with different functional genomic datasets. This specific choice of the epigenomic and sequence grammar feature similarity metric is a result of our investigations on the ENCODE candidate cis-regulatory elements (cCREs) (Moore *et al*., 2020). We identified 103,529 orthologous cCREs between human and mouse using UCSC liftOver (Supplementary Materials Section 1.3) and quantified their epigenomic and sequence grammar feature similarities as described above using the matched ENCODE epigenome datasets and motif scans of the JASPAR database. All the orthogolous cCREs exhibited markedly higher epigenome similarities than randomly matched human and mouse cCREs, supporting a broad level of epigenome conservation between the orthogolous regulatory elements (Fig. 3a). Moreover, promoter like signatures (PLS), proximal enhancer like signatures (pELS), and distal enhancer like signatures (dELS) exhibited monotonically decreasing epigenome conservation which further highlighted the affinity of the epigenomic feature similarity to capture different classes of regulatory elements. This decrease in the epigenomic feature similarity score going from promoters to distal enhancers can be attributed to the rapid divergence of enhancers compared to promoters during regulatory evolution (Cheng *et al*., 2014; Villar *et al*., 2015). Similarly, Fig. 3b illustrates that the sequence grammar similarity captures the degree of sequence conservation across all cCRE categories since UCSC liftOver-defined orthologues are based on pairwise sequence alignment results.

**Figure 3:**
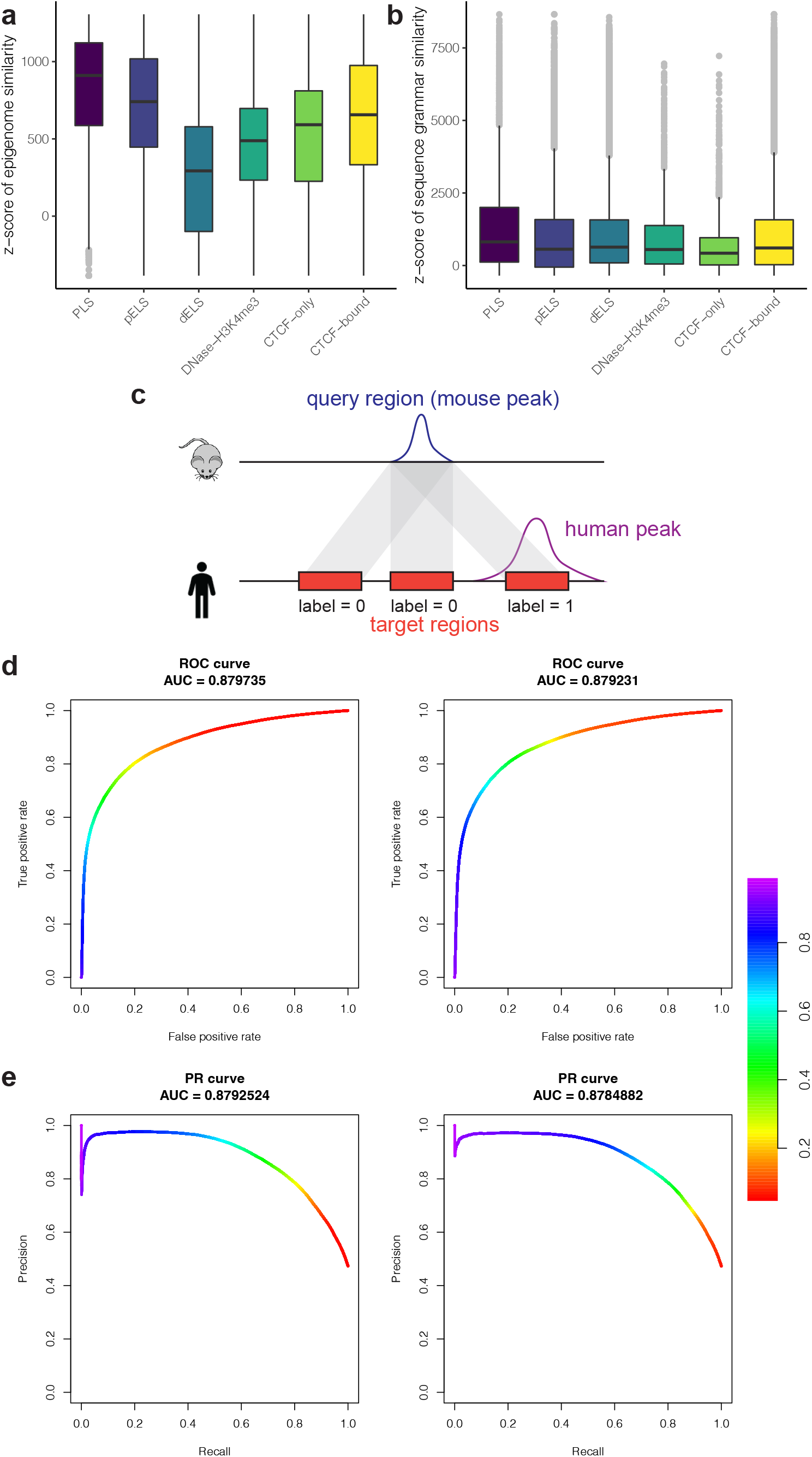
**a-b.** Regulatory information similarity between orthologous cCREs. X-axis: the 6 cCRE categories. A null distribution for each of the similarity scores is estimated by randomly permuting cCREs 10,000 times. Observed similarity scores were transformed into z-scores using the mean and variance estimates of these null distributions. **a.** The z-scores of the epigenomic feature similarity across the 6 cCRE categories. **b.** The z-scores of the sequence grammar similarity across the 6 cCRE categories. **c.** An illustration of labeling candidate target regions of a mouse query region with the corresponding human epigenome peaks. Positive and negative classes are represented by 1 and 0, respectively. The translucent gray bands represent candidate orthologous mappings. **d-e.** The receiver operating characteristic (ROC) and the precision recall (PR) curves with a local window size of 2 kb in the islet ATAC-seq study. Left panel: with the default parameters from the LOOCV experiments; Right panel: with the parameters from the refitted logistic regression. The color denotes the threshold for the logistic probability score.

We also observed that cosine similarity yielded better stability than the Jaccard similarity for binary features which further justified this choice (Supplementary Materials Section 1.3).

#### Parameter tuning with the training module

In order to tune the parameters of AdaLiftOver, we performed leave-one-out cross-validation (LOOCV) with the 67 matched ENCODE epigenome datasets. Specifically, for each fold of the cross-validation, we ensured that the datasets from the same tissue as the validation dataset were excluded from the training set (e.g., when mapping heart H3K4me3 peaks, all other epigenome data from heart were excluded from the epigenomic feature similarity calculations). We applied AdaLiftOver with a grid of window sizes from 0 to 5 kb incremented by 400 bp. After mapping mouse query regions, i.e., peaks, we labeled the candidate target regions in the human genome as positives if they overlapped with the corresponding human epigenome peaks and as negatives otherwise (Fig. 3c). In order to learn the optimal weights of epigenomic and sequence grammar feature similarities, we fitted a logistic regression model with the two similarity features (Supplementary Materials Section 1.5) and computed the area under receiver operating characteristic (ROC) and precision recall (PR) curves for this logistic fit. Optimizing the area under the ROC and PR curves yielded ~2 kb as the optimal window size (AUROC: 0.820±0.0113, AUPR: 0.604±0.0371; Supplementary Materials Section 1.5). We used 2 kb as the size of the local window in generating candidate target regions for the rest of this manuscript. In contrast to the stable local window size, the optimal logistic regression coefficients exhibited larger variability across different folds of the cross-validation. Therefore, we leveraged the averaged coefficient estimates as the weights for the two similarities in the logistic function (Supplementary Materials Section 1.5). To facilitate training with other model organism data, we implemented an AdaLiftOver training module which allows users to estimate the logistic regression coefficients and experiment with thresholds for the logistic probability score.

To further investigate the robustness of the default parameters of AdaLiftOver set by the LOOCV experiments with the ENCODE repertoire, we applied AdaLiftOver on 46,676 mouse pancreatic islet ATAC-seq peaks (Dong *et al*., 2021) with widths between 150 bp and 3 kb as the query regions and performed the following experiment. After generating the candidate target regions at a grid of window sizes, we evaluated them by fitting the logistic regression with labelled data where the candidate target regions overlapping the gold standard human islet ATAC-seq peaks (Greenwald *et al.*, 2019) were labelled as 1 and the rest as 0. First, we observed that the optimal window size of 2 kb from the LOOCV experiments agreed well with the optimal window size in this experiment (Supplementary Figure 5). Next, we scored the candidate target regions generated at local window size of 2 kb with (1) the default parameters from the LOOCV experiments for the logistic regression and (2) parameters from the refitted logistic regression by labeling the candidate target regions as above. Overall, we observed that performance with parameters (#2 above) tuned on this experiment agreed well with the parameters (#1 above) trained with the LOOCV experiments of the 67 ENCODE epigenome datasets (Figs. 3d-e; AUROC: #1 above 0.880, #2 above 0.879; AUPR: #1 above 0.879, #2 above 0.878). This further justified the default parameter setting in AdaLiftOver. All the ROC and PR calculations were conducted with the R package PRROC (Keilwagen *et al*., 2014; Grau *et al*., 2015).

#### Enrichment analysis of mapped regions

To provide support for the mapped regions in the case studies we presented, we asked whether they resided within genomic regions with relevant epigenomic/genic features in the mapped genome more than expected by chance. The null distributions for quantifying these enrichments were adjusted for background genomic factors such as chromosomes and the PhyloP scores of the mapped regions (Supplementary Materials Section 1.8).

## 3 Results

### 3.1 AdaLiftOver generates candidate human epigenome datasets from model organism data

#### Case study I: CD4 and CD8 ATAC-seq peaks from human and mouse

In order to derive human ATAC-seq peaks from the mouse orthologues, Hook and McCallion, 2020 investigated mapping mouse ATAC-seq peak summits using UCSC liftOver or bnMapper and identified an extension of 250 bp on both sides of the summit as performing the best. We first compared AdaLiftOver with this approach by utilizing the same set of human and mouse ATAC-seq summits from the CD4 and CD8 cells (Hook and McCallion, 2020). AdaLiftOver, with default parameters (trained in the LOOCV experiments of the ENCODE repertoire) and at a logistic probability threshold of 0.4, mapped the least number of mouse ATAC-seq peaks (38,779 for CD4; 39,096 for CD8) while attaining the highest number of true positives (23,868 for CD4; 24,208 for CD8), exceeding those of the Hook and McCallion, 2020 (Fig. 4a, Supplementary Materials Section 1.6). As expected, bnMapper and UCSC liftOver behave similarly with an average precision of 47.6% compared to AdaLiftOver’s 61.7%. EpiAlignment yields a high false positive rate due to mapping 87.3% of the regions. In contrast, thresholding the logistic probability score helps AdaLiftOver to remove false positives that might have sequence conservation but lack epigenome conservation and, thus, improve the precision of mapping.

**Figure 4:**
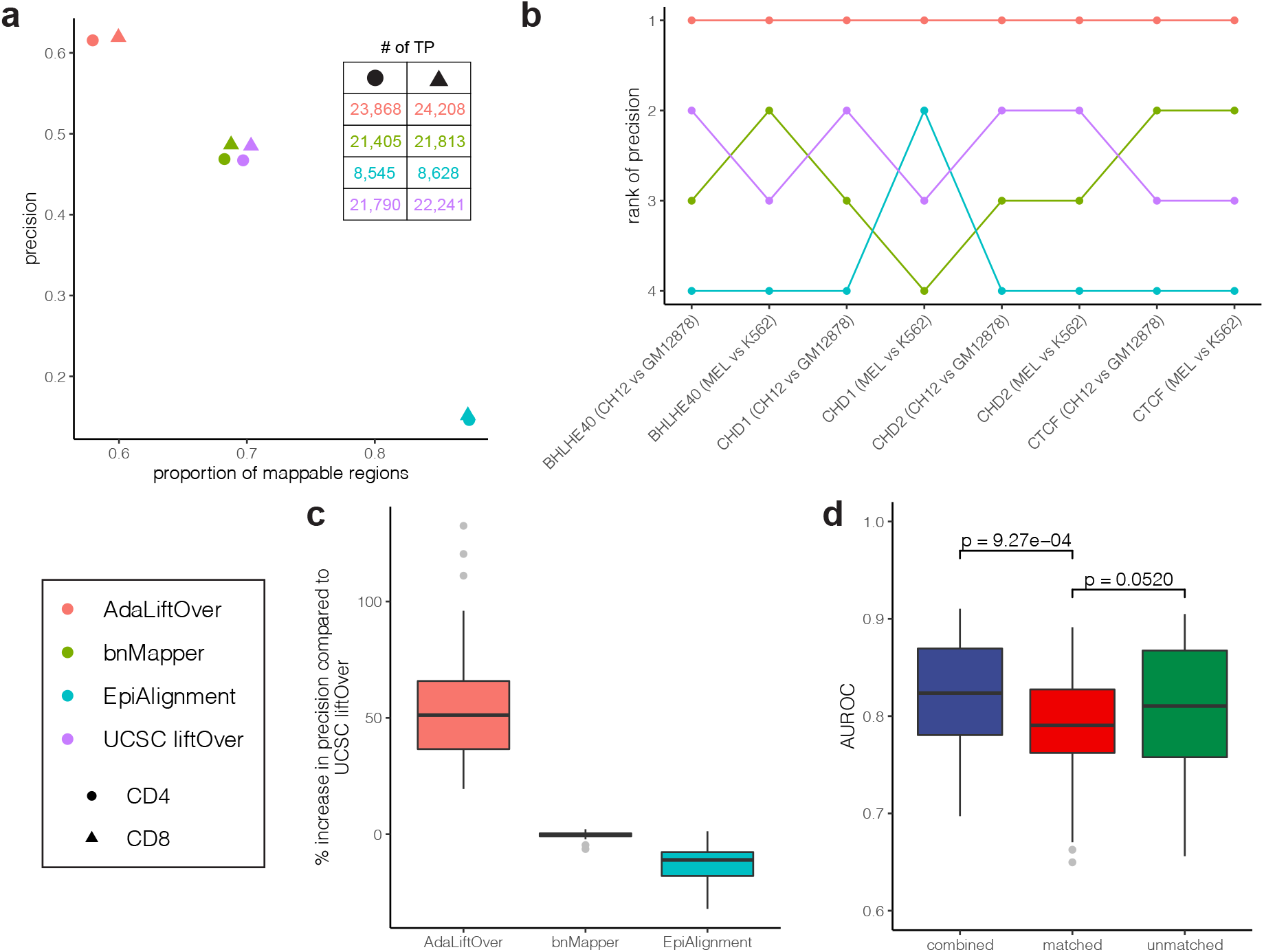
**a.** Comparison of four orthologous mapping methods (color) over two pairs of ATAC-seq datasets (shape). Y-axis: precision is defined as the (# mapped regions with label 1) / (# mapped regions). The actual numbers of true positives for each method are listed in the top-right table. **b.** Comparison of the ranks of the four orthologous mapping methods (color) in terms of their precision over eight pairs of TF ChIP-seq datasets that included results from EpiAlignment. **c.** Percentage increase in precision of the methods compared to the state-of-the-art UCSC liftOver over 55 pairs of TF ChIP-seq datasets. **d.** Comparison of performances of AdaLiftOver under three different configurations of epigenome dataset repertoire. Y-axis: area under the receiver operating characteristic curve. unmatched: the default 67 pairs of ENCODE datasets excluding the relevant cell type; matched: open chromatin regions from the same cell type only; combined: a weighted combination between the previous two where the relevant cell type receives 10x more weight. The p-values are computed from Mann–Whitney U tests.

#### Case study II: Large-scale TF ChIP-seq data from orthologous human and mouse cell lines

We sought to benchmark AdaLiftOver against other orthologous mapping methods for a larger collection of epigenome datasets. We utilized 55 human-mouse orthologous TF ChIP-seq peak sets (Cheng *et al.*, 2014) from erythrold and lymphoblast cells (Supplementary Materials Section 1.7). Specifically, we applied AdaLiftOver to 55 mouse ChIP-seq datasets with an average of 22,679 peaks. Due to the scalability issue of EpiAlignment, we only applied EpiAlignment on 8 pairs of samples displayed in Fig. 4b. AdaLiftOver achieves the best precision while maintaining, on average, 4,456 true positives compared to UCSC liftOver’s 4,313 true positives (Fig. 4b, Supplementary Materials Section 1.7) for all pairs of orthologous datasets. With an average precision of 0.372, AdaLiftOver boosts the precision by >50% compared to UCSC liftOver (Fig. 4c).

Mapping with AdaLiftOver in the above settings did not include any epigenome datasets from the tissues/cell types relevant to the query regions. Next, we asked whether including epigenomic feature from the relevant tissue/cell type impacted the performance. Specifically, we leveraged two pairs of matched ENCODE DNase-seq and ATAC-seq datasets from erythrold and lymphoblast cells (Supplementary Materials Section 1.7). We observed that AdaLiftOver has a better predictive power with the default 67 out-of-sample epigenome datasets than using the relevant open chromation regions alone (Fig. 4d). This demonstrates the practical robustness of AdaLiftOver. As expected, combining all datasets (both the default repertoire and the epigenome dataset from the relevant tissue/cell type) yielded the best performance.

### 3.2 AdaLiftOver enables orthologous mapping for human GWAS SNPs

We considered three sets of GWAS SNPs to evaluate AdaLiftOver and the existing methods. The evaluations are largely based on evaluating whether the mapped regions were enriched in biologically relevant genomic regions (i.e., peaks from epigenome datasets of relevant cell types that were not utilized in mapping, neighbourhood of GWAS phenotype-relevant genes) in the mapped genome.

#### Case study III: Schizophrenia GWAS SNPs

To evaluate the performances of AdaLiftOver and other methods for mapping GWAS SNPs to model organism mouse, we investigated the 1,648 fine-mapped Schizophrenia (SCZ) GWAS SNPs prioritized by Hook and McCallion, 2020. We further utilized the large collection of mouse ATAC-seq data from 25 different brain cell populations out of 6 cell types (Hook and McCallion, 2020) to evaluate the mapping results by their enrichment in the relevant cell populations. UCSC liftOver maps 715 (43.3%) GWAS SNPs where 1.47% ~ 19.1% of them overlap with each of the 25 mouse ATAC-seq datasets. In comparison, AdaLiftOver maps 612 (37.5%) GWAS SNPs and achieves a higher average precision of 8.07%. Fig. 5a illustrates that AdaLiftOver displays a similar trend with stronger enrichment patterns than UCSC liftOver for the relevant cell populations. Consistent with the S-LDSC enrichment results by Hook and McCallion, 2020, we find that SCZ GWAS SNPs are enriched in chromatin accessible regions of all the excitatory neurons and all the inhibitory neurons except PV and VIP (Supplementary Materials Section 1.8). AdaLiftOver largely preserves the biological information of the SCZ GWAS SNPs after cross-species mapping.

**Figure 5:**
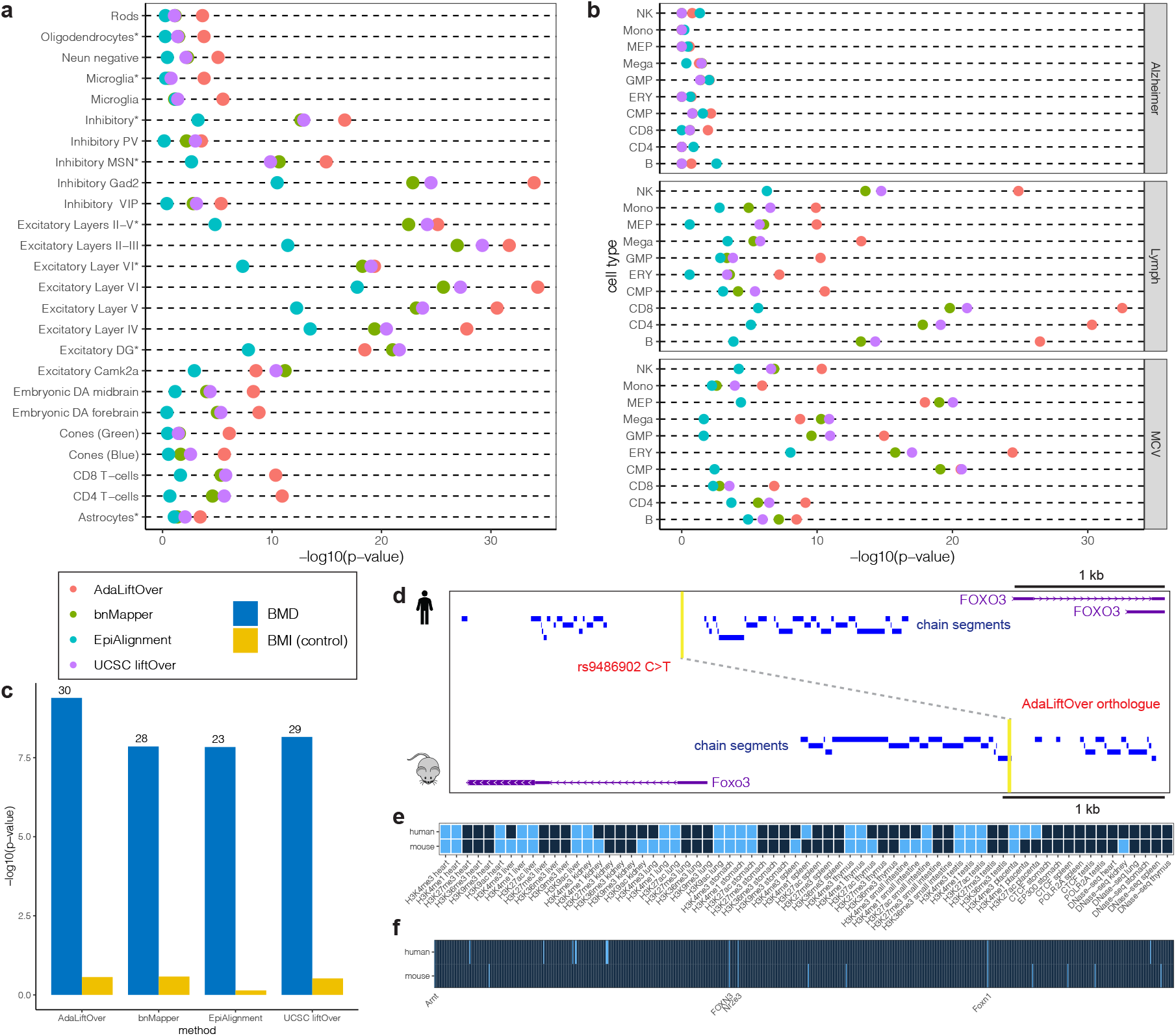
**a.** Enrichment analysis for fine-mapped Schizophrenia GWAS SNPs with respect to ATAC-seq peaks from 25 cell populations. **b.** Enrichment analysis for fine-mapped GWAS SNPs from three traits (Lymph and MCV are hematopoietic traits, Alzheimer is a control trait) with respect to 10 hematopoietic ATAC-seq peaks. Lymph: lymphocyte count; MCV: mean corpuscular volume. **c.** Enrichment analysis for fine-mapped bone mineral density (BMD) GWAS SNPs with respect to 52 mouse BMD genes. BMI: body mass index (control trait). The numbers of BMD genes mapped are labeled on the top of each bar. **d-f.** AdaLiftOver rescues and maps a BMD GWAS rs9486902 at the *FOXO3* promoter in human to *Foxo3* promoter in mouse. *d.* The human GWAS and AdaLiftOver-derived mouse orthologue are highlighted in yellow and linked by a gray dashed line. The GWAS SNP rs9486902 fails to map using UCSC liftOver. *e-f.* The binary epigenomic and sequence grammar feature profiles of the GWAS SNP and its AdaLiftOver orthologue. Light blue and dark blue denotes 1 (overlap) and 0 (not overlap), respectively.

#### Case study IV: Hematopoiesis GWAS SNPs

To further evaluate AdaLiftOver and other methods in the GWAS SNPs setting, we leveraged human fine-mapped GWAS data for four hematopoietic traits: mean corpuscular volume (MCV), mean platelet volume (MPV), monocyte count (Mono), and lymphocyte count (Lymph) (Ulirsch *et al*., 2019). We mapped these SNPs to the mouse genome and performed enrichment analysis with the mouse ATAC-seq peaks from 10 blood cell types (Xiang *et al*., 2020). The enrichment analysis demonstrates that SNPs for MCV are enriched in chromatin accessible regions of ERY, MEP, CMP, and GMP cells; SNPs for Lymph are enriched for NK, CD4, CD8, and B cells (Fig. 5b, Supplementary Materials Section 1.9). These observations are largely consistent with the g-chromVAR (Ulirsch et al., 2019) with the exception of MPV and Mono traits (Supplementary Materials Section 1.9). For these set of mappings, AdaLiftOver and UCSC liftOver perform similarly in terms of enrichments of their mappings in relevant cell types.

#### Case study V: Bone Mineral Density GWAS SNPs

We next showcase how AdaLiftOver can be utilized to map UK Biobank SNPs (Sudlow *et al*., 2015) to mouse for further investigation. In a study of osteoporosis, Swan *et al*., 2020 reported 200 mouse genes as significantly altering bone mineral density (BMD) using BMD measures obtained from a large pool of mice genetically modified for deletion of individual genes. Swan *et al.*, 2020 identified 52 human orthologues of these mouse BMD genes within a 250 kb distance range of UK Biobank BMD GWAS SNPs. To further leverage this knockout mouse resource, we mapped 3,125 fine-mapped BMD GWAS SNPs (Morris *et al*., 2019, the UK Biobank with PIP ≥ 0.1) to the mouse genome and evaluated their enrichment for BMD genes. We used 3,601 body mass index GWAS SNPs from the UK Biobank as negative controls. Compared to other methods, AdaLiftOver achieves the best enrichment results and is capable of identifying the most number of BMD genes (30/52) as relevant for human BMD GWAS SNPs (Fig. 5c, Supplementary Materials Section 1.10). In order to associate more BMD genes with human GWAS SNPs, we then interrogated a larger set of 116,402 GWAS SNPs from the UK Biobank (PIP ≥ 0.001). As a result, AdaLiftOver maps to 90 BMD genes with 65.2% increase in precision compared to UCSC liftOver (Supplementary Materials Section 1.10). Figs. 5d-f illustrate an example where UCSC liftOver does not map any SNPs to the vicinity of the mouse BMD gene *Foxo3* gene but AdaLiftOver is able to rescue *Fox3* with mapping of a BMD GWAS SNP. The SNP rs9486902 resides at the promoter region of human gene *FOXO3* while it is located in a gap among human-mouse chain segments leading to a miss by UCSC liftOver. AdaLiftOver is able to identify a mouse orthologue at the promoter region of *Foxo3* that has similar epigenomic features (Fig. 5e) and sequence grammar (Fig. 5f). In particular, these human and mouse orthologous regions anchored by the *FOXO3* and *Foxo3* genes share common transcription factor binding site motifs that are relevant for BMD. Specifically, ARNT co-binds with Ahr which negatively influences osteoblast proliferation (Yu *et al*., 2014). FOXN3 interacts with Menin, the product of *MEN1,* which influences bone metabolism (Kaji, 2012). Nr2e3, as a nuclear receptor (Oh *et al.*, 2008), is related to human disorders including reduced bone mineral density (Achermann and Jameson, 2003; Achermann et al., 2017). Overall, 25.3% of these UK Biobank (PIP ≥ 0.001) GWAS SNPs can be mapped; however, the majority of them (92.7%) are mapped to “desert” regions that are 250 kb away from the 200 BMD gene promoters, emphasizing the necessity for follow-up with 3D genome profiling assays such as pcHi-C (Mifsud *et al*., 2015) and its variants.

## 4 Discussion

We developed AdaLiftOver to enable mapping of noncoding regions between human and model organism genomes. AdaLiftOver goes beyond traditional sequence alignment of comparative genomics for lifting over between genomes and simultaneously incorporates comparative epigenomics and sequence grammar similarity. To the best of our knowledge, this is the first systematic benchmark study of different orthologous mapping methods with comprehensive real biological data applications. Compared to other methods, AdaLiftOver is more accurate and robust, and offers a computationally inexpensive way of generating hard-to-obtain functional genomic datasets in other genomes by incorporating epigenomic and sequence grammar features. Table. 1 further summarizes the flexibility, scalability, and running time of AdaLiftOver compared to existing methods.

**Table 1:** Technical comparison among existing orthologues mapping methods. Sequence alignment: the dependence on the UCSC sequence alignment framework. Running time: the average running time for ~20,000 TF ChIP-seq peaks. Generalizability: whether or not the method supports any genomes.

| method | sequence alignment | epigenomic features | sequence grammar features | scalability | running time | generalizability |
| --- | --- | --- | --- | --- | --- | --- |
| **AdaLiftOver** | Yes | Multiple | Yes | Yes | ~20 minutes | Yes |
| **bnMapper (Denas *et al.*, 2015)** | Yes | No | No | Yes | ~10 minutes | Yes |
| **EpiAlignment (Lu *et al.*, 2019)** | Yes | One | No | No | ~3 hours | No |
| **UCSC liftOver (Hinrichs *et al.*, 2006)** | Yes | No | No | Yes | ~10 seconds | Yes |

We found that the majority of orthologues of GWAS SNPs tend to have an enriched but low overlapping percentage with related open chromatin regions in mouse. We expect this result to improve as the epigenomic features leveraged span more cell types and dynamic conditions. In particular, developmental and disease trajectories revealed by single cell ATAC-seq might provide more enrichment for orthologues of GWAS SNPs. With a more comprehensive epigenome space, AdaLiftOver can serve as a versatile approach for pinpointing potential GWAS orthologues in a model organism and can facilitate high-throughput perturbation experiments. Currently, AdaLiftOver is restricted to binary features due to the space requirements and time complexity. We expect a more computationally efficient implementation of AdaLiftOver to incorporate features at other scales.

## References

Achermann, J. C. and Jameson, J. L. (2003). Human disorders caused by nuclear receptor gene mutations. Pure and applied chemistry, 75(11-12), 1785–1796.

Achermann, J. C., Schwabe, J., Fairall, L., Chatterjee, K., et al. (2017). Genetic disorders of nuclear receptors. The Journal of Clinical Investigation, 127(4), 1181–1192.

Alipanahi, B., Delong, A., Weirauch, M. T., and Frey, B. J. (2015). Predicting the sequence specificities of dna-and rna-binding proteins by deep learning. Nature biotechnology, 33(8), 831–838.

Brawand, D., Soumillon, M., Necsulea, A., Julien, P., Csárdi, G., Harrigan, P., Weier, M., Liechti, A., Aximu-Petri, A., Kircher, M., et al. (2011). The evolution of gene expression levels in mammalian organs. Nature, 478(7369), 343–348.

Castro-Mondragon, J. A., Riudavets-Puig, R., Rauluseviciute, I., Berhanu Lemma, R., Turchi, L., Blanc-Mathieu, R., Lucas, J., Boddie, P., Khan, A., Manosalva Pérez, N., et al. (2022). Jaspar 2022: the 9th release of the open-access database of transcription factor binding profiles. Nucleic acids research, 50(D1), D165–D173.

Cheng, Y., Ma, Z., Kim, B.-H., Wu, W., Cayting, P., Boyle, A. P., Sundaram, V., Xing, X., Dogan, N., Li, J., et al. (2014). Principles of regulatory information conservation between mouse and human. Nature, 515(7527), 371–375.

Denas, O., Sandstrom, R., Cheng, Y., Beal, K., Herrero, J., Hardison, R. C., and Taylor, J. (2015). Genome-wide comparative analysis reveals human-mouse regulatory landscape and evolution. BMC genomics, 16(1), 1–9.

Dong, C., Simonett, S. P., Shin, S., Stapleton, D. S., Schueler, K. L., Churchill, G. A., Lu, L., Liu, X., Jin, F., Li, Y., et al. (2021). Infima leverages multi-omics model organism data to identify effector genes of human gwas variants. Genome biology, 22(1), 1–32.

Earl, D., Nguyen, N., Hickey, G., Harris, R. S., Fitzgerald, S., Beal, K., Seledtsov, I., Molodtsov, V., Raney, B. J., Clawson, H., et al. (2014). Alignathon: a competitive assessment of wholegenome alignment methods. Genome research, 24(12), 2077–2089.

Gallagher, M. D. and Chen-Plotkin, A. S. (2018). The post-gwas era: from association to function. The American Journal of Human Genetics, 102(5), 717–730.

Gjoneska, E., Pfenning, A. R., Mathys, H., Quon, G., Kundaje, A., Tsai, L.-H., and Kellis, M. (2015). Conserved epigenomic signals in mice and humans reveal immune basis of alzheimer’s disease. Nature, 518(7539), 365–369.

Grau, J., Grosse, I., and Keilwagen, J. (2015). Prroc: computing and visualizing precision-recall and receiver operating characteristic curves in r. Bioinformatics, 31(15), 2595–2597.

Greenwald, W. W., Chiou, J., Yan, J., Qiu, Y., Dai, N., Wang, A., Nariai, N., Aylward, A., Han, J. Y., Kadakia, N., et al. (2019). Pancreatic islet chromatin accessibility and conformation reveals distal enhancer networks of type 2 diabetes risk. Nature communications, 10(1), 1–12.

Hinrichs, A. S., Karolchik, D., Baertsch, R., Barber, G. P., Bejerano, G., Clawson, H., Diekhans, M., Furey, T. S., Harte, R. A., Hsu, F., et al. (2006). The ucsc genome browser database: update 2006. Nucleic acids research, 34(suppl_1), D590–D598.

Hook, P. W. and McCallion, A. S. (2020). Leveraging mouse chromatin data for heritability enrichment informs common disease architecture and reveals cortical layer contributions to schizophrenia. Genome research, 30(4), 528–539.

Kaji, H. (2012). Menin and bone metabolism. Journal of bone and mineral metabolism, 30(4), 381–387.

Keilwagen, J., Grosse, I., and Grau, J. (2014). Area under precision-recall curves for weighted and unweighted data. PloS one, 9(3), e92209.

Kelley, D. R. (2020). Cross-species regulatory sequence activity prediction. PLoS computational biology, 16(7), e1008050.

Kelley, D. R., Snoek, J., and Rinn, J. L. (2016). Basset: learning the regulatory code of the accessible genome with deep convolutional neural networks. Genome research, 26(7), 990–999.

Kelley, D. R., Reshef, Y. A., Bileschi, M., Belanger, D., McLean, C. Y., and Snoek, J. (2018). Sequential regulatory activity prediction across chromosomes with convolutional neural networks. Genome research, 28(5), 739–750.

Kwon, S. B. and Ernst, J. (2021). Learning a genome-wide score of human–mouse conservation at the functional genomics level. Nature communications, 12(1), 1–14.

Lu, J., Cao, X., and Zhong, S. (2019). Epialignment: alignment with both dna sequence and epigenomic data. Nucleic acids research, 47(W1), W11–W19.

Mifsud, B., Tavares-Cadete, F., Young, A. N., Sugar, R., Schoenfelder, S., Ferreira, L., Wingett, S. W., Andrews, S., Grey, W., Ewels, P. A., et al. (2015). Mapping long-range promoter contacts in human cells with high-resolution capture hi-c. Nature genetics, 47(6), 598–606.

Minnoye, L., Taskiran, I. I., Mauduit, D., Fazio, M., Van Aerschot, L., Hulselmans, G., Christiaens, V., Makhzami, S., Seltenhammer, M., Karras, P., et al. (2020). Cross-species analysis of enhancer logic using deep learning. Genome research, 30(12), 1815–1834.

Moore, J. E., Purcaro, M. J., Pratt, H. E., Epstein, C. B., Shoresh, N., Adrian, J., Kawli, T., Davis, C. A., Dobin, A., Kaul, R., et al. (2020). Expanded encyclopaedias of dna elements in the human and mouse genomes. Nature, 583(7818), 699–710.

Morris, J. A., Kemp, J. P., Youlten, S. E., Laurent, L., Logan, J. G., Chai, R. C., Vulpescu, N. A., Forgetta, V., Kleinman, A., Mohanty, S. T., et al. (2019). An atlas of genetic influences on osteoporosis in humans and mice. Nature genetics, 51(2), 258–266.

Odom, D. T., Dowell, R. D., Jacobsen, E. S., Gordon, W., Danford, T. W., MacIsaac, K. D., Rolfe, P. A., Conboy, C. M., Gifford, D. K., and Fraenkel, E. (2007). Tissue-specific transcriptional regulation has diverged significantly between human and mouse. Nature genetics, 39(6), 730–732.

Oh, E. C., Cheng, H., Hao, H., Jia, L., Khan, N. W., and Swaroop, A. (2008). Rod differentiation factor nrl activates the expression of nuclear receptor nr2e3 to suppress the development of cone photoreceptors. Brain research, 1236, 16–29.

Schep, A. N., Wu, B., Buenrostro, J. D., and Greenleaf, W. J. (2017). chromvar: inferring transcription-factor-associated accessibility from single-cell epigenomic data. Nature methods, 14(10), 975–978.

Sudlow, C., Gallacher, J., Allen, N., Beral, V., Burton, P., Danesh, J., Downey, P., Elliott, P., Green, J., Landray, M., et al. (2015). Uk biobank: an open access resource for identifying the causes of a wide range of complex diseases of middle and old age. PLoS medicine, 12(3), e1001779.

Swan, A. L., Schütt, C., Rozman, J., del Mar Muñiz Moreno, M., Brandmaier, S., Simon, M., Leuchtenberger, S., Griffiths, M., Brommage, R., Keskivali-Bond, P., et al. (2020). Mouse mutant phenotyping at scale reveals novel genes controlling bone mineral density. PLoS genetics, 16(12), e1009190.

Ulirsch, J. C., Lareau, C. A., Bao, E. L., Ludwig, L. S., Guo, M. H., Benner, C., Satpathy, A. T., Kartha, V. K., Salem, R. M., Hirschhorn, J. N., et al. (2019). Interrogation of human hematopoiesis at single-cell and single-variant resolution. Nature genetics, 51(4), 683–693.

Vierstra, J., Rynes, E., Sandstrom, R., Zhang, M., Canfield, T., Hansen, R. S., Stehling-Sun, S., Sabo, P. J., Byron, R., Humbert, R., et al. (2014). Mouse regulatory dna landscapes reveal global principles of cis-regulatory evolution. Science, 346(6212), 1007–1012.

Villar, D., Berthelot, C., Aldridge, S., Rayner, T. F., Lukk, M., Pignatelli, M., Park, T. J., Deaville, R., Erichsen, J. T., Jasinska, A. J., et al. (2015). Enhancer evolution across 20 mammalian species. Cell, 160(3), 554–566.

Welter, D., MacArthur, J., Morales, J., Burdett, T., Hall, P., Junkins, H., Klemm, A., Flicek, P., Manolio, T., Hindorff, L., et al. (2014). The nhgri gwas catalog, a curated resource of snp-trait associations. Nucleic acids research, 42(D1), D1001–D1006.

Xiang, G., Keller, C. A., Heuston, E., Giardine, B. M., An, L., Wixom, A. Q., Miller, A., Cockburn, A., Sauria, M. E., Weaver, K., et al. (2020). An integrative view of the regulatory and transcriptional landscapes in mouse hematopoiesis. Genome research, 30(3), 472–484.

Yu, H., Du, Y., Zhang, X., Sun, Y., Li, S., Dou, Y., Li, Z., Yuan, H., and Zhao, W. (2014). The aryl hydrocarbon receptor suppresses osteoblast proliferation and differentiation through the activation of the erk signaling pathway. Toxicology and applied pharmacology, 280(3), 502–510.

Zhou, J. and Troyanskaya, O. G. (2015). Predicting effects of noncoding variants with deep learning-based sequence model. Nature methods, 12(10), 931–934.

Zhou, J., Theesfeld, C. L., Yao, K., Chen, K. M., Wong, A. K., and Troyanskaya, O. G. (2018). Deep learning sequence-based ab initio prediction of variant effects on expression and disease risk. Nature genetics, 50(8), 1171–1179.

