## Supplementary Materials for "AdaLiftOver: High-resolution identification of orthologous regulatory elements with adaptive liftOver"

Chenyang Dong[1] and Sündüz Keleş[1,2,*]

[1]Department of Statistics,
[2]Department of Biostatistics and Medical Informatics,

**2 Supplementary Tables** **7**

**3 Supplementary Figures** **23**

### 1 Supplementary Notes

#### 1.1 The orthologous gene promoters

We downloaded NCBI RefSeq gene annotation files from `https://hgdownload.soe.ucsc.edu/goldenPath/hg38/bigZips/genes/hg38.ncbiRefSeq.gtf.gz` and `https://hgdownload.soe.ucsc.edu/goldenPath/mm10/bigZips/genes/mm10.ncbiRefSeq.gtf.gz`. We utilized the human-mouse homology table from the Jackson Laboratory `http://www.informatics.jax.org/downloads/reports/HOM_MouseHumanSequence.rpt` and identified 16,374 orthologous genes in total. We defined the promoter region of a gene as 2 kb upstream and 200 bp downstream of the transcription start site (TSS). We used the reciprocal chain files "hg38.mm10.rbest.chain" and "mm10.hg38.rbest.chain" to determine the mappings between the orthologous promoters. We first utilized the UCSC liftOver (The R function "rtracklayer::liftOver()") to map all human/mouse promoters to the mouse/human genome and overlapped them with the corresponding counterparts. When a human/mouse promoter failed to map, we extended the promoters by 2 kb on both sides and remapped the extended region. In this context, we defined the positive and negative mappings as follows:

- True positive (TP): a promoter maps and overlaps with the correct counterpart.
- False positive (FP): a promoter maps but does not overlap with the correct counterpart.
- True negative (TN): a promoter fails to map or overlap with the correct counterpart after extending by 2 kb on both sides.
- False negative (FN): a promoter fails to map but it maps to the correct counterpart after extending by 2 kb on both sides.

The results are summarized in Supplementary Table 1.

#### 1.2 The ENCODE datasets in the AdaLiftOver default repertoire for computing epigenomic feature similarities

The complete list of the datasets is provided in Supplementary Table 2.

#### 1.3 Evaluating similarity metrics with the ENCODE cCRE data

The orthologous ENCODE cCRE data (v3) from both hg38 and mm10 were downloaded from `https://screen.encodeproject.org/`. We utilized the R function "rtracklayer::liftOver()" with the UCSC chain file "hg38.mm10.rbest.chain" to map human cCREs to mouse. The 103,529 orthologues were determined by requiring an overlap of at least 50% of the bases between the human cCRE orthologues and mouse cCREs. The human cCRE categories before and after mapping are summarized in Supplementary Figure 1.

With the 103,529 orthologous cCREs, we further evaluated the combinations of three different similarity metrics (Pearson correlation, cosine similarity, and soft cosine similarity[1]) and three different data formats summarizing the epigenomic signals in the form of peaks (BED format), fold change of normalized ChIP reads over control reads (Fold change BigWig), and p-values comparing ChIP and control reads along the genome (P-value BigWig). We specifically considered:

[1] soft_cosine$(u, v) = \frac{\sum_{i,j} s_{ij} u_i v_j}{\sqrt{\sum_{i,j} s_{ij} u_i u_j}\sqrt{\sum_{i,j} s_{ij} v_i v_j}}$, where $s_{ij}$ was estimated by the sample Pearson correlation between the $i$-th and $j$-th entries of epigenomic feature vectors for all human and mouse cCREs.

- BED format: we overlapped each cCRE with the list of peaks and quantified the strength of the signal as the percentage overlap with the peaks (numerical value between 0 and 1).
- Fold change BigWig format: we computed the arithmetic average of BigWig signals for each cCRE.
- P-value BigWig format: we computed the geometric average of BigWig signals for each cCRE.

Supplementary Figure 2 illustrates that the soft cosine similarity underperforms compared to the other metrics, and the fold-change BigWig format is outperformed by the p-value BigWig format as benchmarked by the z-scores. We observe that the BED format and the p-value BigWig format yield the best performance when combined with the Pearson correlation and the cosine similarity, respectively. Although the p-value BigWig format (with the Pearson correlation metric) achieves higher average z-scores compared to the BED format (with the cosine similarity metric), it exhibits larger variability across all 6 cCRE categories (Supplementary Figure 2). Therefore, the numerical features derived from the BED format data are as robust and informative as the BigWig formatted data.

The binary features constructed by overlapping the cCREs with the peaks (BED format data) attain similar performance as the numerical features derived from the BED format data (with the cosine similarity metric) (Supplementary Figure 3). A comparison of Supplementary Figures 2 and 3 indicates that the cosine similarity, which shows less variability than the Jaccard similarity, is a better similarity metric for binary vectors. We also incorporated the Jaccard similarity option in AdaLiftOver.

##### 1.4 Discussion on the choice of data formats and similarity metrics.

*Epigenomic feature similarity.* The 67 pairs of BigWig files from the ENCODE repertoire required $\sim$10 GB disk storage after being compressed from the $\sim$150 GB raw data to the Run-Length-Encoding structure separated by chromosomes. In contrast, only $<$50 MB space was required to store the binary signals of the BED files from the same ENCODE repertoire. Querying the BigWig format data, while it theoretically takes O(1) time complexity, was computational prohibitive for small servers and personal laptops. For a scientific computing server with Intel Xeon CPU E5-2680 v4 2.40GHz, it took $\sim$20 minutes to load the data in R and $\sim$24 hours to handle $\sim$100,000 orthologous genomic regions. However, querying the BED format data only took $\sim$1 minute for the same task. Further memory issues arose as the number of parallel jobs increase, rendering the overall approach using the BigWig numeric signals infeasible. On the contrary, the BED binary signals scaled well on a regular laptop. Therefore, AdaLiftOver leveraged binary features derived from the BED files with the cosine similarity metric for optimizing the disk storage, the computational speed, the scalability, and the performance robustness (Supplementary Figures 2 and 3).

*Sequence grammar feature similarity.* The pre-computed occurrence data for the 841 JASPAR motifs in the human and mouse genomes occupied $\sim$200 GB disk storage. Due to the high space requirement and to enable applicability to any model organism genome, we chose the motifmatchr R package, accelerated by C++ libraries, to perform on-demand motif scanning using PFM/PWMs. After quantifying motif occurrences, we used binary features and cosine similarity metric for the sequence grammar.

#### 1.5 Parameter tuning for AdaLiftOver

*Leave-one-out cross validation (LOOCV) with the ENCODE datasets.* For each of the 67 ENCODE epigenome datasets in Supplementary Table 2, we utilized the remaining ENCODE datasets excluding those from the same tissue as the training data to compute epigenomic feature similarities. For the candidate target regions in human, we overlapped them with the corresponding human peaks and labeled each pair of orthologues. With a grid of window sizes, Supplementary Figure 4 illustrates that the optimal window size lies between 1 kb and 3 kb by considering the trade-off between AUPR and AUROC. In practice, we suggest using a window size within this range.

At window = 2 kb, we summarize below the estimated logistic regression coefficients (mean $\pm$ s.e.; $\hat{\beta}_0, \hat{\beta}_1, \hat{\beta}_2$ denote the estimated coefficients for the intercept, epigenomic feature similarity, and sequence grammar feature similarity, respectively):

- $\hat{\beta}_0 = -4.16 \pm 0.231$
- $\hat{\beta}_1 = 5.40 \pm 0.349$
- $\hat{\beta}_2 = 2.06 \pm 0.212$

We used these set of parameters as default for the rest of the manuscript. With the constraint of filtering at most 50% of candidate target regions, we identified logistic probability score thresholds 0.376$\pm$0.0374 that maximize the precision 0.554$\pm$0.0371 using the AdaLiftOver training module. We selected the score threshold = 0.4 for the case studies on ATAC-seq and ChIP-seq peaks. The effects of selecting a logistic probability score threshold can be visually investigated with ROC/PR curves. The users may increase/decrease the score threshold if more stringent/lenient results are needed. We also considered an interaction term between the epigenomic feature similarity and the sequence grammar feature similarity in the logistic regression. However, the area under the curves did not yield significant differences with or without an interaction term (Mann-Whitney test p-values: 0.9752 for AUROC, 0.9539 for AUPR). AdaLiftOver implementation allows inclusion of such an interaction term which might impact the results in other datasets.

#### 1.6 The CD4 and CD8 ATAC-seq data case

We applied the existing orthologous mapping methods to 66,921 mouse CD4 and 65,209 mouse CD8 ATAC-seq peak summits from Hook and McCallion, 2020. We further examined the choice of the logistic probability score threshold (Supplementary Figure 7).

The settings of orthologous mapping methods:

- AdaLiftOver: threshold = 0.4. The chain file is “mm10.hg38.rbest.chain”
- bnMapper: Default parameters. The chain file is “mm10.hg38.rbest.chain”
- UCSC liftOver: The R function “rtracklayer::liftOver()”. The chain file is “mm10ToHg38.over.chain”
- EpiAlignment: window = 2 kb. We used paired H3K4me3 ChIP-seq datasets from Adult B-lymphocytes as the orthologous epigenomic features.

Then, we extended the mouse-derived summits to 501 bp and overlapped the resulting peaks with 40,530 CD4 ATAC-seq peaks and 40,916 CD8 ATAC-seq peaks in human. The results are summarized in Supplementary Table 3, which are better than the number of overlapped peaks (true positives) reported in Hook and McCallion, 2020.

#### 1.7 ENCODE and mouse ENCODE TF ChIP-seq datasets

We utilized 55 pairs of TF ChIP-seq datasets from Cheng *et al.*, 2014 (Supplementary Table 4). The methods were run under the following settings:

- AdaLiftOver: threshold = 0.4. The chain file is "mm10.hg38.rbest.chain"
- bnMapper: Default parameters. The chain file is "mm10.hg38.rbest.chain"
- UCSC liftOver: The R function "rtracklayer::liftOver()". The chain file is "mm10ToHg38.over.chain"
- EpiAlignment: window = 2 kb. In order to tune EpiAlignment fairly, for MEL vs K562 samples, we used paired Erythrold cells H3K4me3 ChIP-seq data as the epigenome data input; For CH12 vs GM12878 samples, we used Lymphoblast cells H3K4me3 ChIP-seq data as the epigenome data input.

The results are summarized in in Supplementary Table 5.

We then downloaded the following ENCODE ATAC-seq/DNase-seq datasets to evaluate the performances of AdaLiftOver with matched tissues/cell types:

- mouse-CH12, DNase-seq, ENCFF855RCO.bed
- human-GM12878, ATAC-seq, ENCFF470YYO.bed
- mouse-MEL, DNase-seq, ENCFF726EZS.bed
- human-K562, ATAC-seq, ENCFF558BLC.bed

When combining the above epigenome profiles with the default ENCODE epigenome repertoire, we set the weights ($w$) of the epigenomic features from the matched tissues by a factor of 10 more in the similarity calculations.

#### 1.8 Schizophrenia GWAS SNPs

We utilized 1,648 SCZ GWAS SNPs (annotation PIP $\geq$ 0.1) and 25 mouse ATAC-seq datasets from Hook and McCallion, 2020. The benchmarking results are summarized in Supplementary Table 6.

The methods were run under the following settings:

- AdaLiftOver: threshold = 0.2. The chain file is "hg19.mm10.rbest.chain"
- bnMapper: Default parameters. The chain file is "hg19.mm10.rbest.chain"
- UCSC liftOver: The R function "rtracklayer::liftOver()". The chain file is "hg19ToMm10.over.chain"
- EpiAlignment: window = 2 kb. GWAS SNPs from hg19 were converted to hg38 with UCSC liftOver using the chain file "hg19ToHg38.over.chain". Due to the lack of BMD related orthologous epigenome data, we used orthologous cCRE data as the user inputted epigenomic signals in order to tune EpiAlignment fairly.

To provide support for the mapped GWAS SNPs, we asked whether ATAC-seq peaks of relevant cell types overlapped with the mapped GWAS SNPs more than expected by chance. Our calculations took into account background genomic characteristics such as sequence conservation as quantified by the PhyloP scores[2] and genomic coordinates as quantified by the

[2]The PhyloP dataset was downloaded from `http://hgdownload.cse.ucsc.edu/goldenpath/mm10/phyloP60way/mm10.60way.phyloP60way.bw`

chromosomes that the ATAC-seq peaks reside in. We first binned all the open chromatin regions[3] of the genome into intervals of 100 bp and stratified these with respect to their PhyloP scores discretized at 0.1 resolution for each chromosome separately. Then, we computed the empirical probability of overlap ($p_i$) with the mapped GWAS SNPs within each strata $S_i$, $i = 1 \cdots, I$, where index $i$ runs over the PhyloP score grid and the numbers of chromosomes. Next, for a given ATAC-seq peak set of size $P$, we first calculated the expected proportion of overlap with the GWAS SNPs as $p_0 = \frac{1}{P}\sum_p \sum_i I(p\text{-th peak is in strata } i)p_i$. Using this null proportion, we evaluated the enrichment with a Binomial test that assessed whether the observed overlap of the ATAC-seq peak set with the mapped GWAS SNPs is larger than that can be explained by this expected proportion of overlap.

##### 1.9 Hematopoiesis GWAS SNPs

We leveraged 10 blood-related mouse ATAC-seq peaks (Xiang *et al.*, 2020) and fine-mapped hematopoietic GWAS SNPs (Ulirsch *et al.*, 2019) with PIP $\geq$ 0.1 from 4 blood traits: MCV, MPV, Mono, Lymph. The settings of the methods compared and the enrichment analysis were the same as the SCZ GWAS case. The benchmarking results are summarized in Supplementary Table 7. The enrichment results for Mono (monocyte count) and MPV (mean platelet volume) are displayed in Supplementary Figure 8.

##### 1.10 Bone Mineral Density GWAS SNPs

The promoter regions of BMD genes were defined as 2 kb upstream and 200 bp downstream of TSS. We utilized 1,097 fine-mapped BMD GWAS SNPs from Morris *et al.*, 2019 as well as 2,088 estimated BMD fine-mapped GWAS SNPs from the UK Biobank (SuSIE PIP $\geq$ 0.1). For negative controls, we utilized a comparable number of 3,601 BMI SNPs from the UK Biobank (SuSIE PIP $\geq$ 0.05). The settings of the methods compared and the enrichment analysis were the same as the SCZ GWAS case. The regions for enrichment analysis were the 52 BMD gene promoters and the constructed universe consisted of all orthologous gene promoters identified in Section 1.1. The results are summarized in Supplementary Table 8.

We further used 116,402 fine-mapped GWAS SNPs from the UK Biobank (SuSIE PIP $\geq$ 0.001). We didn't run EpiAlignment due to the lack of scalability of this method. The results are summarized in Supplementary Table 9.

[3]The 100bp-binned open chromatin regions as the union of all target ATAC-seq peaks extended by 1 kb.

#### 2 Supplementary Tables

| | TP | FP | TN | FN |
|---|---|---|---|---|
| **human to mouse** | 12,760 | 2,813 | 592 | 209 |
| **mouse to human** | 12,760 | 2,525 | 933 | 156 |

Supplementary Table 1: Mapping of the 16,374 orthologous gene promoters between human and mouse.

| | Assay | Target | Tissue | mouse file id | human file id |
|---|---|---|---|---|---|
| 1 | ChIP-seq | H3K4me3 | heart | ENCFF925JLJ.bed | ENCFF685WYG.bed |
| 2 | ChIP-seq | H3K4me1 | heart | ENCFF577SAP.bed | ENCFF109AVD.bed |
| 3 | ChIP-seq | H3K27me3 | heart | ENCFF337GMV.bed | ENCFF594SIJ.bed |
| 4 | ChIP-seq | H3K36me3 | heart | ENCFF412PEF.bed | ENCFF501WXA.bed |
| 5 | ChIP-seq | H3K9me3 | heart | ENCFF018ZTP.bed | ENCFF472HHB.bed |
| 6 | ChIP-seq | H3K9ac | heart | ENCFF043BXJ.bed | ENCFF305FQY.bed |
| 7 | ChIP-seq | H3K4me3 | liver | ENCFF233NKJ.bed | ENCFF178DYP.bed |
| 8 | ChIP-seq | H3K4me1 | liver | ENCFF626UMI.bed | ENCFF872NIB.bed |
| 9 | ChIP-seq | H3K27ac | liver | ENCFF734MGU.bed | ENCFF805YRQ.bed |
| 10 | ChIP-seq | H3K27me3 | liver | ENCFF290NCY.bed | ENCFF977LFS.bed |
| 11 | ChIP-seq | H3K36me3 | liver | ENCFF958ZWB.bed | ENCFF050ODT.bed |
| 12 | ChIP-seq | H3K9me3 | liver | ENCFF763XZB.bed | ENCFF093NPF.bed |
| 13 | ChIP-seq | H3K9ac | liver | ENCFF025NBX.bed | ENCFF766JJU.bed |
| 14 | ChIP-seq | H3K4me3 | kidney | ENCFF870HWW.bed | ENCFF917SIV.bed |
| 15 | ChIP-seq | H3K4me1 | kidney | ENCFF661JHH.bed | ENCFF224BUW.bed |
| 16 | ChIP-seq | H3K27me3 | kidney | ENCFF357TDA.bed | ENCFF775YUI.bed |
| 17 | ChIP-seq | H3K36me3 | kidney | ENCFF219HDC.bed | ENCFF839LUB.bed |
| 18 | ChIP-seq | H3K9me3 | kidney | ENCFF616DMW.bed | ENCFF163PII.bed |
| 19 | ChIP-seq | H3K9ac | kidney | ENCFF226CKB.bed | ENCFF138PYK.bed |
| 20 | ChIP-seq | H3K4me3 | lung | ENCFF142XFR.bed | ENCFF254OZF.bed |
| 21 | ChIP-seq | H3K4me1 | lung | ENCFF262JML.bed | ENCFF908EUN.bed |
| 22 | ChIP-seq | H3K27ac | lung | ENCFF455ADY.bed | ENCFF314QTE.bed |
| 23 | ChIP-seq | H3K27me3 | lung | ENCFF536OYS.bed | ENCFF112CDW.bed |
| 24 | ChIP-seq | H3K36me3 | lung | ENCFF262QMM.bed | ENCFF089HKW.bed |
| 25 | ChIP-seq | H3K9me3 | lung | ENCFF737BJL.bed | ENCFF284QKU.bed |
| 26 | ChIP-seq | H3K9ac | lung | ENCFF871DGI.bed | ENCFF330TSX.bed |
| 27 | ChIP-seq | H3K4me3 | stomach | ENCFF902SVE.bed | ENCFF494BWU.bed |
| 28 | ChIP-seq | H3K4me1 | stomach | ENCFF377USJ.bed | ENCFF475EJU.bed |
| 29 | ChIP-seq | H3K27ac | stomach | ENCFF643VAF.bed | ENCFF978AOD.bed |
| 30 | ChIP-seq | H3K27me3 | stomach | ENCFF993YFO.bed | ENCFF692HXY.bed |
| 31 | ChIP-seq | H3K36me3 | stomach | ENCFF704KRU.bed | ENCFF243EAW.bed |
| 32 | ChIP-seq | H3K9me3 | stomach | ENCFF401WHJ.bed | ENCFF299EII.bed |
| 33 | ChIP-seq | H3K4me3 | spleen | ENCFF040ATH.bed | ENCFF636LHJ.bed |
| 34 | ChIP-seq | H3K4me1 | spleen | ENCFF332TVZ.bed | ENCFF997KKX.bed |
| 35 | ChIP-seq | H3K27ac | spleen | ENCFF675SAZ.bed | ENCFF805FFP.bed |
| 36 | ChIP-seq | H3K27me3 | spleen | ENCFF472CRY.bed | ENCFF867NGE.bed |
| 37 | ChIP-seq | H3K36me3 | spleen | ENCFF962GUW.bed | ENCFF593PUF.bed |
| 38 | ChIP-seq | H3K4me3 | thymus | ENCFF462HWJ.bed | ENCFF963LSO.bed |
| 39 | ChIP-seq | H3K4me1 | thymus | ENCFF149ZSO.bed | ENCFF821ZRD.bed |

Continued on next page

Supplementary Table 2 – continued from previous page

| | Assay | Target | Tissue | mouse file id | human file id |
|---|---|---|---|---|---|
| 40 | ChIP-seq | H3K27ac | thymus | ENCFF476TDY.bed | ENCFF668BZJ.bed |
| 41 | ChIP-seq | H3K27me3 | thymus | ENCFF497BVT.bed | ENCFF641CCT.bed |
| 42 | ChIP-seq | H3K36me3 | thymus | ENCFF394EHO.bed | ENCFF869FKJ.bed |
| 43 | ChIP-seq | H3K4me3 | small intestine | ENCFF250OKJ.bed | ENCFF638SMB.bed |
| 44 | ChIP-seq | H3K4me1 | small intestine | ENCFF061UOC.bed | ENCFF377UAM.bed |
| 45 | ChIP-seq | H3K27ac | small intestine | ENCFF371CEP.bed | ENCFF074WDV.bed |
| 46 | ChIP-seq | H3K27me3 | small intestine | ENCFF359SCM.bed | ENCFF644QDF.bed |
| 47 | ChIP-seq | H3K36me3 | small intestine | ENCFF841SYQ.bed | ENCFF694AKL.bed |
| 48 | ChIP-seq | H3K4me3 | testis | ENCFF582PBS.bed | ENCFF244NRV.bed |
| 49 | ChIP-seq | H3K4me1 | testis | ENCFF868HGX.bed | ENCFF600MJO.bed |
| 50 | ChIP-seq | H3K27ac | testis | ENCFF047JTI.bed | ENCFF540YUQ.bed |
| 51 | ChIP-seq | H3K27me3 | testis | ENCFF031KAC.bed | ENCFF319CMK.bed |
| 52 | ChIP-seq | H3K36me3 | testis | ENCFF187TKX.bed | ENCFF469ASG.bed |
| 53 | ChIP-seq | H3K4me3 | placenta | ENCFF296GDX.bed | ENCFF829GCX.bed |
| 54 | ChIP-seq | H3K4me1 | placenta | ENCFF499ASE.bed | ENCFF867CRG.bed |
| 55 | ChIP-seq | H3K27ac | placenta | ENCFF226XOZ.bed | ENCFF297NSU.bed |
| 56 | ChIP-seq | CTCF | stomach | ENCFF907XEG.bed | ENCFF017NAF.bed |
| 57 | ChIP-seq | EP300 | stomach | ENCFF132DDY.bed | ENCFF099PQM.bed |
| 58 | ChIP-seq | CTCF | spleen | ENCFF089EWX.bed | ENCFF645YCX.bed |
| 59 | ChIP-seq | POLR2A | spleen | ENCFF313QNI.bed | ENCFF683YQZ.bed |
| 60 | ChIP-seq | CTCF | testis | ENCFF033UQX.bed | ENCFF917CYG.bed |
| 61 | ChIP-seq | POLR2A | testis | ENCFF789JUX.bed | ENCFF967TFR.bed |
| 62 | DNase-seq | | heart | ENCFF846XYQ.bed | ENCFF861CQD.bed |
| 63 | DNase-seq | | kidney | ENCFF890GCF.bed | ENCFF388NNQ.bed |
| 64 | DNase-seq | | lung | ENCFF459KBR.bed | ENCFF691GQY.bed |
| 65 | DNase-seq | | stomach | ENCFF220GXC.bed | ENCFF731YGM.bed |
| 66 | DNase-seq | | spleen | ENCFF343KXE.bed | ENCFF523NCT.bed |
| 67 | DNase-seq | | thymus | ENCFF266KGB.bed | ENCFF494PNZ.bed |

Supplementary Table 2: The 67 matched ENCODE datasets between mouse and human. These datasets make up AdaLiftOver's default epigenome repertoire.

| | # peaks | # mapped peaks | # overlapped peaks | precision | sample | method |
|---|---|---|---|---|---|---|
| 1 | 66921 | 38779 | 23868 | 0.62 | CD4 | AdaLiftOver |
| 2 | 66921 | 45670 | 21405 | 0.47 | CD4 | bnMapper |
| 3 | 66921 | 58472 | 8545 | 0.15 | CD4 | EpiAlignment |
| 4 | 66921 | 46646 | 21790 | 0.47 | CD4 | UCSC liftOver |
| 5 | 65209 | 39096 | 24208 | 0.62 | CD8 | AdaLiftOver |
| 6 | 65209 | 44840 | 21813 | 0.49 | CD8 | bnMapper |
| 7 | 65209 | 56911 | 8628 | 0.15 | CD8 | EpiAlignment |
| 8 | 65209 | 45857 | 22241 | 0.49 | CD8 | UCSC liftOver |

Supplementary Table 3: Benchmarking AdaLiftOver on ATAC-seq peaks from CD4 and CD8 cells.

| | TF | human-K562 | mouse-MEL | human-GM12878 | mouse-CH12 |
|---|---|---|---|---|---|
| 1 | BHLHE40 | Yes | Yes | Yes | Yes |
| 2 | CHD1 | Yes | Yes | Yes | Yes |
| 3 | CHD2 | Yes | Yes | Yes | Yes |
| 4 | CTCF | Yes | Yes | Yes | Yes |
| 5 | E2F4 | Yes | Yes | Yes | Yes |
| 6 | ELF1 | Yes | Yes | Yes | Yes |
| 7 | EP300 | Yes | Yes | Yes | Yes |
| 8 | ETS1 | Yes | Yes | Yes | Yes |
| 9 | GATA1 | Yes | Yes | No | No |
| 10 | JUND | Yes | Yes | Yes | Yes |
| 11 | MAFK | Yes | Yes | No | No |
| 12 | MAX | Yes | Yes | Yes | Yes |
| 13 | MAZ | Yes | Yes | Yes | Yes |
| 14 | MEF2A | Yes | Yes | Yes | Yes |
| 15 | MXI1 | Yes | Yes | Yes | Yes |
| 16 | MYC | Yes | Yes | Yes | Yes |
| 17 | NRF1 | Yes | Yes | Yes | Yes |
| 18 | POLR2A | Yes | Yes | Yes | Yes |
| 19 | RAD21 | Yes | Yes | Yes | Yes |
| 20 | RCOR1 | Yes | Yes | Yes | Yes |
| 21 | RDBP | Yes | Yes | No | No |
| 22 | SIN3A | Yes | Yes | Yes | Yes |
| 23 | SMC3 | Yes | Yes | Yes | Yes |
| 24 | TAL1 | Yes | Yes | No | No |
| 25 | TBP | Yes | Yes | Yes | Yes |
| 26 | UBTF | Yes | Yes | No | No |
| 27 | USF1 | Yes | Yes | Yes | Yes |
| 28 | USF2 | Yes | Yes | Yes | Yes |
| 29 | IRF4 | No | No | Yes | Yes |
| 30 | KAT2A | No | No | Yes | Yes |
| 31 | PAX5 | No | No | Yes | Yes |
| 32 | TCF12 | No | No | Yes | Yes |

Supplementary Table 4: Orthologous human and mouse TF ChIP-seq datasets used for benchmarking.

| | # peaks | # map | # overlap | precision | sample | method |
|---|---|---|---|---|---|---|
| 1 | 43498 | 18834 | 6761 | 0.36 | BHLHE40 (CH12 vs GM12878) | AdaLiftOver |
| 2 | 43498 | 32606 | 5970 | 0.18 | BHLHE40 (CH12 vs GM12878) | bnMapper |
| 3 | 43498 | 37751 | 6059 | 0.16 | BHLHE40 (CH12 vs GM12878) | EpiAlignment |
| 4 | 43498 | 33589 | 6152 | 0.18 | BHLHE40 (CH12 vs GM12878) | UCSC liftOver |
| 5 | 16325 | 8080 | 2911 | 0.36 | BHLHE40 (MEL vs K562) | AdaLiftOver |
| 6 | 16325 | 11716 | 2450 | 0.21 | BHLHE40 (MEL vs K562) | bnMapper |
| 7 | 16325 | 13803 | 2471 | 0.18 | BHLHE40 (MEL vs K562) | EpiAlignment |
| 8 | 16325 | 12076 | 2518 | 0.21 | BHLHE40 (MEL vs K562) | UCSC liftOver |
| 9 | 11286 | 6984 | 1417 | 0.20 | CHD1 (CH12 vs GM12878) | AdaLiftOver |
| 10 | 11286 | 9209 | 1049 | 0.11 | CHD1 (CH12 vs GM12878) | bnMapper |
| 11 | 11286 | 10071 | 1118 | 0.11 | CHD1 (CH12 vs GM12878) | EpiAlignment |
| 12 | 11286 | 9640 | 1114 | 0.12 | CHD1 (CH12 vs GM12878) | UCSC liftOver |
| 13 | 8170 | 6103 | 2269 | 0.37 | CHD1 (MEL vs K562) | AdaLiftOver |
| 14 | 8170 | 6686 | 1566 | 0.23 | CHD1 (MEL vs K562) | bnMapper |
| 15 | 8170 | 7336 | 1774 | 0.24 | CHD1 (MEL vs K562) | EpiAlignment |
| 16 | 8170 | 7036 | 1680 | 0.24 | CHD1 (MEL vs K562) | UCSC liftOver |
| 17 | 19553 | 11302 | 5176 | 0.46 | CHD2 (CH12 vs GM12878) | AdaLiftOver |
| 18 | 19553 | 15579 | 4589 | 0.29 | CHD2 (CH12 vs GM12878) | bnMapper |
| 19 | 19553 | 17297 | 4633 | 0.27 | CHD2 (CH12 vs GM12878) | EpiAlignment |
| 20 | 19553 | 16135 | 4787 | 0.30 | CHD2 (CH12 vs GM12878) | UCSC liftOver |
| 21 | 5062 | 3497 | 1727 | 0.49 | CHD2 (MEL vs K562) | AdaLiftOver |
| 22 | 5062 | 4009 | 1542 | 0.38 | CHD2 (MEL vs K562) | bnMapper |
| 23 | 5062 | 4428 | 1553 | 0.35 | CHD2 (MEL vs K562) | EpiAlignment |
| 24 | 5062 | 4194 | 1615 | 0.39 | CHD2 (MEL vs K562) | UCSC liftOver |
| 25 | 60828 | 22812 | 13086 | 0.57 | CTCF (CH12 vs GM12878) | AdaLiftOver |
| 26 | 60828 | 36477 | 14918 | 0.41 | CTCF (CH12 vs GM12878) | bnMapper |
| 27 | 60828 | 48665 | 13944 | 0.29 | CTCF (CH12 vs GM12878) | EpiAlignment |
| 28 | 60828 | 37944 | 15395 | 0.41 | CTCF (CH12 vs GM12878) | UCSC liftOver |
| 29 | 46608 | 18792 | 12015 | 0.64 | CTCF (MEL vs K562) | AdaLiftOver |
| 30 | 46608 | 26427 | 13862 | 0.52 | CTCF (MEL vs K562) | bnMapper |
| 31 | 46608 | 36707 | 12972 | 0.35 | CTCF (MEL vs K562) | EpiAlignment |
| 32 | 46608 | 27488 | 14299 | 0.52 | CTCF (MEL vs K562) | UCSC liftOver |
| 33 | 5579 | 3863 | 1358 | 0.35 | E2F4 (CH12 vs GM12878) | AdaLiftOver |
| 34 | 5579 | 4477 | 1140 | 0.25 | E2F4 (CH12 vs GM12878) | bnMapper |
| 35 | 5579 | 4693 | 1207 | 0.26 | E2F4 (CH12 vs GM12878) | UCSC liftOver |
| 36 | 4122 | 3319 | 1813 | 0.55 | E2F4 (MEL vs K562) | AdaLiftOver |
| 37 | 4122 | 3362 | 1509 | 0.45 | E2F4 (MEL vs K562) | bnMapper |
| 38 | 4122 | 3540 | 1590 | 0.45 | E2F4 (MEL vs K562) | UCSC liftOver |
| 39 | 26680 | 16032 | 9681 | 0.60 | ELF1 (CH12 vs GM12878) | AdaLiftOver |
| 40 | 26680 | 20849 | 8384 | 0.40 | ELF1 (CH12 vs GM12878) | bnMapper |
| 41 | 26680 | 21699 | 8775 | 0.40 | ELF1 (CH12 vs GM12878) | UCSC liftOver |
| 42 | 18949 | 12227 | 7511 | 0.61 | ELF1 (MEL vs K562) | AdaLiftOver |
| 43 | 18949 | 14536 | 6310 | 0.43 | ELF1 (MEL vs K562) | bnMapper |
| 44 | 18949 | 15086 | 6568 | 0.44 | ELF1 (MEL vs K562) | UCSC liftOver |
| 45 | 33645 | 10803 | 1858 | 0.17 | EP300 (CH12 vs GM12878) | AdaLiftOver |
| 46 | 33645 | 24466 | 2364 | 0.10 | EP300 (CH12 vs GM12878) | bnMapper |
| 47 | 33645 | 25215 | 2411 | 0.10 | EP300 (CH12 vs GM12878) | UCSC liftOver |
| 48 | 41335 | 14609 | 3117 | 0.21 | EP300 (MEL vs K562) | AdaLiftOver |
| 49 | 41335 | 27145 | 3028 | 0.11 | EP300 (MEL vs K562) | bnMapper |

Continued on next page

**Supplementary Table 5 – continued from previous page**

| | # peaks | # map | # overlap | precision | sample | method |
|---|---|---|---|---|---|---|
| 50 | 41335 | 28088 | 3120 | 0.11 | EP300 (MEL vs K562) | UCSC liftOver |
| 51 | 29200 | 11825 | 3068 | 0.26 | ETS1 (CH12 vs GM12878) | AdaLiftOver |
| 52 | 29200 | 21951 | 2540 | 0.12 | ETS1 (CH12 vs GM12878) | bnMapper |
| 53 | 29200 | 22658 | 2667 | 0.12 | ETS1 (CH12 vs GM12878) | UCSC liftOver |
| 54 | 38734 | 16858 | 4641 | 0.28 | ETS1 (MEL vs K562) | AdaLiftOver |
| 55 | 38734 | 26613 | 3440 | 0.13 | ETS1 (MEL vs K562) | bnMapper |
| 56 | 38734 | 27550 | 3593 | 0.13 | ETS1 (MEL vs K562) | UCSC liftOver |
| 57 | 45252 | 14359 | 1146 | 0.08 | GATA1 (MEL vs K562) | AdaLiftOver |
| 58 | 45252 | 28062 | 1413 | 0.05 | GATA1 (MEL vs K562) | bnMapper |
| 59 | 45252 | 28968 | 1428 | 0.05 | GATA1 (MEL vs K562) | UCSC liftOver |
| 60 | 41363 | 18562 | 3853 | 0.21 | IRF4 (CH12 vs GM12878) | AdaLiftOver |
| 61 | 41363 | 31104 | 3918 | 0.13 | IRF4 (CH12 vs GM12878) | bnMapper |
| 62 | 41363 | 32142 | 4040 | 0.13 | IRF4 (CH12 vs GM12878) | UCSC liftOver |
| 63 | 15762 | 5550 | 618 | 0.11 | JUND (CH12 vs GM12878) | AdaLiftOver |
| 64 | 15762 | 11411 | 750 | 0.07 | JUND (CH12 vs GM12878) | bnMapper |
| 65 | 15762 | 11786 | 765 | 0.06 | JUND (CH12 vs GM12878) | UCSC liftOver |
| 66 | 6674 | 2818 | 1073 | 0.38 | JUND (MEL vs K562) | AdaLiftOver |
| 67 | 6674 | 4355 | 988 | 0.23 | JUND (MEL vs K562) | bnMapper |
| 68 | 6674 | 4510 | 1017 | 0.23 | JUND (MEL vs K562) | UCSC liftOver |
| 69 | 10569 | 7685 | 74 | 0.01 | KAT2A (CH12 vs GM12878) | AdaLiftOver |
| 70 | 10569 | 8726 | 60 | 0.01 | KAT2A (CH12 vs GM12878) | bnMapper |
| 71 | 10569 | 9152 | 63 | 0.01 | KAT2A (CH12 vs GM12878) | UCSC liftOver |
| 72 | 2351 | 943 | 281 | 0.30 | MAFK (MEL vs K562) | AdaLiftOver |
| 73 | 2351 | 1479 | 328 | 0.22 | MAFK (MEL vs K562) | bnMapper |
| 74 | 2351 | 1525 | 333 | 0.22 | MAFK (MEL vs K562) | UCSC liftOver |
| 75 | 27057 | 15319 | 5860 | 0.38 | MAX (CH12 vs GM12878) | AdaLiftOver |
| 76 | 27057 | 20988 | 4783 | 0.23 | MAX (CH12 vs GM12878) | bnMapper |
| 77 | 27057 | 21865 | 5023 | 0.23 | MAX (CH12 vs GM12878) | UCSC liftOver |
| 78 | 24511 | 14527 | 8186 | 0.56 | MAX (MEL vs K562) | AdaLiftOver |
| 79 | 24511 | 18210 | 6446 | 0.35 | MAX (MEL vs K562) | bnMapper |
| 80 | 24511 | 19038 | 6787 | 0.36 | MAX (MEL vs K562) | UCSC liftOver |
| 81 | 17739 | 11632 | 6426 | 0.55 | MAZ (CH12 vs GM12878) | AdaLiftOver |
| 82 | 17739 | 14709 | 5622 | 0.38 | MAZ (CH12 vs GM12878) | bnMapper |
| 83 | 17739 | 15291 | 5899 | 0.39 | MAZ (CH12 vs GM12878) | UCSC liftOver |
| 84 | 16851 | 11707 | 7695 | 0.66 | MAZ (MEL vs K562) | AdaLiftOver |
| 85 | 16851 | 13589 | 6631 | 0.49 | MAZ (MEL vs K562) | bnMapper |
| 86 | 16851 | 14166 | 6949 | 0.49 | MAZ (MEL vs K562) | UCSC liftOver |
| 87 | 28178 | 13764 | 3342 | 0.24 | MEF2A (CH12 vs GM12878) | AdaLiftOver |
| 88 | 28178 | 21854 | 3511 | 0.16 | MEF2A (CH12 vs GM12878) | bnMapper |
| 89 | 28178 | 22638 | 3637 | 0.16 | MEF2A (CH12 vs GM12878) | UCSC liftOver |
| 90 | 4620 | 2065 | 294 | 0.14 | MEF2A (MEL vs K562) | AdaLiftOver |
| 91 | 4620 | 3361 | 301 | 0.09 | MEF2A (MEL vs K562) | bnMapper |
| 92 | 4620 | 3467 | 306 | 0.09 | MEF2A (MEL vs K562) | UCSC liftOver |
| 93 | 23755 | 16096 | 8317 | 0.52 | MXI1 (CH12 vs GM12878) | AdaLiftOver |
| 94 | 23755 | 19488 | 6960 | 0.36 | MXI1 (CH12 vs GM12878) | bnMapper |
| 95 | 23755 | 20445 | 7400 | 0.36 | MXI1 (CH12 vs GM12878) | UCSC liftOver |
| 96 | 29509 | 18473 | 4588 | 0.25 | MXI1 (MEL vs K562) | AdaLiftOver |
| 97 | 29509 | 22747 | 3529 | 0.16 | MXI1 (MEL vs K562) | bnMapper |

Continued on next page

Supplementary Table 5 – continued from previous page

| | # peaks | # map | # overlap | precision | sample | method |
|---|---|---|---|---|---|---|
| 98 | 29509 | 23869 | 3786 | 0.16 | MXI1 (MEL vs K562) | UCSC liftOver |
| 99 | 26233 | 14407 | 2517 | 0.17 | MYC (CH12 vs GM12878) | AdaLiftOver |
| 100 | 26233 | 19896 | 1860 | 0.09 | MYC (CH12 vs GM12878) | bnMapper |
| 101 | 26233 | 20648 | 1967 | 0.10 | MYC (CH12 vs GM12878) | UCSC liftOver |
| 102 | 23675 | 13283 | 7219 | 0.54 | MYC (MEL vs K562) | AdaLiftOver |
| 103 | 23675 | 17136 | 5759 | 0.34 | MYC (MEL vs K562) | bnMapper |
| 104 | 23675 | 17897 | 6055 | 0.34 | MYC (MEL vs K562) | UCSC liftOver |
| 105 | 16163 | 11913 | 3667 | 0.31 | NRF1 (CH12 vs GM12878) | AdaLiftOver |
| 106 | 16163 | 13206 | 3173 | 0.24 | NRF1 (CH12 vs GM12878) | bnMapper |
| 107 | 16163 | 13823 | 3354 | 0.24 | NRF1 (CH12 vs GM12878) | UCSC liftOver |
| 108 | 10599 | 8591 | 2700 | 0.31 | NRF1 (MEL vs K562) | AdaLiftOver |
| 109 | 10599 | 8861 | 2309 | 0.26 | NRF1 (MEL vs K562) | bnMapper |
| 110 | 10599 | 9254 | 2435 | 0.26 | NRF1 (MEL vs K562) | UCSC liftOver |
| 111 | 194 | 58 | 23 | 0.40 | PAX5 (CH12 vs GM12878) | AdaLiftOver |
| 112 | 194 | 119 | 19 | 0.16 | PAX5 (CH12 vs GM12878) | bnMapper |
| 113 | 194 | 129 | 22 | 0.17 | PAX5 (CH12 vs GM12878) | UCSC liftOver |
| 114 | 23696 | 16595 | 10486 | 0.63 | POLR2A (CH12 vs GM12878) | AdaLiftOver |
| 115 | 23696 | 18700 | 9080 | 0.49 | POLR2A (CH12 vs GM12878) | bnMapper |
| 116 | 23696 | 19889 | 9771 | 0.49 | POLR2A (CH12 vs GM12878) | UCSC liftOver |
| 117 | 21898 | 16012 | 3375 | 0.21 | POLR2A (MEL vs K562) | AdaLiftOver |
| 118 | 21898 | 16949 | 2761 | 0.16 | POLR2A (MEL vs K562) | bnMapper |
| 119 | 21898 | 18123 | 2943 | 0.16 | POLR2A (MEL vs K562) | UCSC liftOver |
| 120 | 41545 | 16323 | 10413 | 0.64 | RAD21 (CH12 vs GM12878) | AdaLiftOver |
| 121 | 41545 | 26278 | 12384 | 0.47 | RAD21 (CH12 vs GM12878) | bnMapper |
| 122 | 41545 | 27311 | 12736 | 0.47 | RAD21 (CH12 vs GM12878) | UCSC liftOver |
| 123 | 30908 | 14131 | 7430 | 0.53 | RAD21 (MEL vs K562) | AdaLiftOver |
| 124 | 30908 | 19912 | 8258 | 0.41 | RAD21 (MEL vs K562) | bnMapper |
| 125 | 30908 | 20695 | 8470 | 0.41 | RAD21 (MEL vs K562) | UCSC liftOver |
| 126 | 10893 | 5326 | 918 | 0.17 | RCOR1 (CH12 vs GM12878) | AdaLiftOver |
| 127 | 10893 | 8549 | 893 | 0.10 | RCOR1 (CH12 vs GM12878) | bnMapper |
| 128 | 10893 | 8877 | 929 | 0.10 | RCOR1 (CH12 vs GM12878) | UCSC liftOver |
| 129 | 25999 | 9929 | 678 | 0.07 | RCOR1 (MEL vs K562) | AdaLiftOver |
| 130 | 25999 | 17522 | 663 | 0.04 | RCOR1 (MEL vs K562) | bnMapper |
| 131 | 25999 | 18154 | 685 | 0.04 | RCOR1 (MEL vs K562) | UCSC liftOver |
| 132 | 17422 | 13091 | 397 | 0.03 | RDBP (MEL vs K562) | AdaLiftOver |
| 133 | 17422 | 14093 | 311 | 0.02 | RDBP (MEL vs K562) | bnMapper |
| 134 | 17422 | 14906 | 345 | 0.02 | RDBP (MEL vs K562) | UCSC liftOver |
| 135 | 27103 | 17821 | 7015 | 0.39 | SIN3A (CH12 vs GM12878) | AdaLiftOver |
| 136 | 27103 | 22261 | 5693 | 0.26 | SIN3A (CH12 vs GM12878) | bnMapper |
| 137 | 27103 | 23341 | 6077 | 0.26 | SIN3A (CH12 vs GM12878) | UCSC liftOver |
| 138 | 37908 | 22270 | 8181 | 0.37 | SIN3A (MEL vs K562) | AdaLiftOver |
| 139 | 37908 | 28786 | 6175 | 0.21 | SIN3A (MEL vs K562) | bnMapper |
| 140 | 37908 | 30118 | 6583 | 0.22 | SIN3A (MEL vs K562) | UCSC liftOver |
| 141 | 42669 | 17756 | 10953 | 0.62 | SMC3 (CH12 vs GM12878) | AdaLiftOver |
| 142 | 42669 | 28251 | 12299 | 0.44 | SMC3 (CH12 vs GM12878) | bnMapper |
| 143 | 42669 | 29363 | 12679 | 0.43 | SMC3 (CH12 vs GM12878) | UCSC liftOver |
| 144 | 39493 | 16863 | 9277 | 0.55 | SMC3 (MEL vs K562) | AdaLiftOver |
| 145 | 39493 | 23913 | 9838 | 0.41 | SMC3 (MEL vs K562) | bnMapper |

Continued on next page

**Supplementary Table 5 – continued from previous page**

| | # peaks | # map | # overlap | precision | sample | method |
|---|---|---|---|---|---|---|
| 146 | 39493 | 24866 | 10116 | 0.41 | SMC3 (MEL vs K562) | UCSC liftOver |
| 147 | 18204 | 6135 | 1045 | 0.17 | TAL1 (MEL vs K562) | AdaLiftOver |
| 148 | 18204 | 11767 | 1412 | 0.12 | TAL1 (MEL vs K562) | bnMapper |
| 149 | 18204 | 12163 | 1431 | 0.12 | TAL1 (MEL vs K562) | UCSC liftOver |
| 150 | 20458 | 12056 | 5799 | 0.48 | TBP (CH12 vs GM12878) | AdaLiftOver |
| 151 | 20458 | 15237 | 4950 | 0.32 | TBP (CH12 vs GM12878) | bnMapper |
| 152 | 20458 | 15969 | 5222 | 0.33 | TBP (CH12 vs GM12878) | UCSC liftOver |
| 153 | 26567 | 15495 | 7807 | 0.50 | TBP (MEL vs K562) | AdaLiftOver |
| 154 | 26567 | 18412 | 6140 | 0.33 | TBP (MEL vs K562) | bnMapper |
| 155 | 26567 | 19244 | 6447 | 0.34 | TBP (MEL vs K562) | UCSC liftOver |
| 156 | 32474 | 18158 | 6907 | 0.38 | TCF12 (CH12 vs GM12878) | AdaLiftOver |
| 157 | 32474 | 25471 | 6162 | 0.24 | TCF12 (CH12 vs GM12878) | bnMapper |
| 158 | 32474 | 26585 | 6476 | 0.24 | TCF12 (CH12 vs GM12878) | UCSC liftOver |
| 159 | 5111 | 4406 | 1811 | 0.41 | UBTF (MEL vs K562) | AdaLiftOver |
| 160 | 5111 | 4436 | 1470 | 0.33 | UBTF (MEL vs K562) | bnMapper |
| 161 | 5111 | 4691 | 1562 | 0.33 | UBTF (MEL vs K562) | UCSC liftOver |
| 162 | 8086 | 4116 | 1691 | 0.41 | USF1 (CH12 vs GM12878) | AdaLiftOver |
| 163 | 8086 | 6028 | 1612 | 0.27 | USF1 (CH12 vs GM12878) | bnMapper |
| 164 | 8086 | 6221 | 1660 | 0.27 | USF1 (CH12 vs GM12878) | UCSC liftOver |
| 165 | 19341 | 9449 | 3071 | 0.33 | USF1 (MEL vs K562) | AdaLiftOver |
| 166 | 19341 | 13293 | 2733 | 0.21 | USF1 (MEL vs K562) | bnMapper |
| 167 | 19341 | 13795 | 2838 | 0.21 | USF1 (MEL vs K562) | UCSC liftOver |
| 168 | 5662 | 2822 | 1036 | 0.37 | USF2 (CH12 vs GM12878) | AdaLiftOver |
| 169 | 5662 | 4266 | 981 | 0.23 | USF2 (CH12 vs GM12878) | bnMapper |
| 170 | 5662 | 4389 | 1010 | 0.23 | USF2 (CH12 vs GM12878) | UCSC liftOver |
| 171 | 1310 | 863 | 514 | 0.60 | USF2 (MEL vs K562) | AdaLiftOver |
| 172 | 1310 | 1030 | 504 | 0.49 | USF2 (MEL vs K562) | bnMapper |
| 173 | 1310 | 1062 | 520 | 0.49 | USF2 (MEL vs K562) | UCSC liftOver |

Supplementary Table 5: Benchmarking AdaLiftOver with the TF ChIP-seq datasets. Precision is computed as # of overlap divided by # of mapped.

| | # map | # overlap | precision | method | sample |
|---|---|---|---|---|---|
| 1 | 612 | 16 | 0.03 | AdaLiftOver | Microglia |
| 2 | 612 | 12 | 0.02 | AdaLiftOver | Astrocytes* |
| 3 | 612 | 13 | 0.02 | AdaLiftOver | Cones (Blue) |
| 4 | 612 | 23 | 0.04 | AdaLiftOver | CD4 T-cells |
| 5 | 612 | 22 | 0.04 | AdaLiftOver | CD8 T-cells |
| 6 | 612 | 57 | 0.09 | AdaLiftOver | Excitatory Camk2a |
| 7 | 612 | 12 | 0.02 | AdaLiftOver | Inhibitory PV |
| 8 | 612 | 16 | 0.03 | AdaLiftOver | Inhibitory VIP |
| 9 | 612 | 105 | 0.17 | AdaLiftOver | Excitatory Layers II-III |
| 10 | 612 | 69 | 0.11 | AdaLiftOver | Excitatory Layer VI* |
| 11 | 612 | 97 | 0.16 | AdaLiftOver | Excitatory DG* |
| 12 | 612 | 105 | 0.17 | AdaLiftOver | Excitatory Layers II-V* |
| 13 | 612 | 21 | 0.03 | AdaLiftOver | Embryonic DA forebrain |
| 14 | 612 | 113 | 0.18 | AdaLiftOver | Inhibitory Gad2 |
| 15 | 612 | 14 | 0.02 | AdaLiftOver | Cones (Green) |
| 16 | 612 | 64 | 0.10 | AdaLiftOver | Inhibitory* |
| 17 | 612 | 73 | 0.12 | AdaLiftOver | Inhibitory MSN* |
| 18 | 612 | 20 | 0.03 | AdaLiftOver | Embryonic DA midbrain |
| 19 | 612 | 12 | 0.02 | AdaLiftOver | Microglia* |
| 20 | 612 | 17 | 0.03 | AdaLiftOver | Neun negative |
| 21 | 612 | 117 | 0.19 | AdaLiftOver | Excitatory Layer VI |
| 22 | 612 | 11 | 0.02 | AdaLiftOver | Oligodendrocytes* |
| 23 | 612 | 113 | 0.18 | AdaLiftOver | Excitatory Layer V |
| 24 | 612 | 9 | 0.01 | AdaLiftOver | Rods |
| 25 | 612 | 103 | 0.17 | AdaLiftOver | Excitatory Layer IV |
| 26 | 681 | 8 | 0.01 | bnMapper | Microglia |
| 27 | 681 | 8 | 0.01 | bnMapper | Astrocytes* |
| 28 | 681 | 7 | 0.01 | bnMapper | Cones (Blue) |
| 29 | 681 | 14 | 0.02 | bnMapper | CD4 T-cells |
| 30 | 681 | 15 | 0.02 | bnMapper | CD8 T-cells |
| 31 | 681 | 71 | 0.10 | bnMapper | Excitatory Camk2a |
| 32 | 681 | 10 | 0.01 | bnMapper | Inhibitory PV |
| 33 | 681 | 12 | 0.02 | bnMapper | Inhibitory VIP |
| 34 | 681 | 102 | 0.15 | bnMapper | Excitatory Layers II-III |
| 35 | 681 | 75 | 0.11 | bnMapper | Excitatory Layer VI* |
| 36 | 681 | 108 | 0.16 | bnMapper | Excitatory DG* |
| 37 | 681 | 108 | 0.16 | bnMapper | Excitatory Layers II-V* |
| 38 | 681 | 16 | 0.02 | bnMapper | Embryonic DA forebrain |
| 39 | 681 | 97 | 0.14 | bnMapper | Inhibitory Gad2 |
| 40 | 681 | 7 | 0.01 | bnMapper | Cones (Green) |
| 41 | 681 | 57 | 0.08 | bnMapper | Inhibitory* |
| 42 | 681 | 66 | 0.10 | bnMapper | Inhibitory MSN* |
| 43 | 681 | 14 | 0.02 | bnMapper | Embryonic DA midbrain |
| 44 | 681 | 5 | 0.01 | bnMapper | Microglia* |
| 45 | 681 | 12 | 0.02 | bnMapper | Neun negative |
| 46 | 681 | 107 | 0.16 | bnMapper | Excitatory Layer VI |
| 47 | 681 | 7 | 0.01 | bnMapper | Oligodendrocytes* |
| 48 | 681 | 105 | 0.15 | bnMapper | Excitatory Layer V |
| 49 | 681 | 5 | 0.01 | bnMapper | Rods |

Continued on next page

**Supplementary Table 6 – continued from previous page**

| | # map | # overlap | precision | method | sample |
|---|---|---|---|---|---|
| 50 | 681 | 94 | 0.14 | bnMapper | Excitatory Layer IV |
| 51 | 1209 | 9 | 0.01 | EpiAlignment | Microglia |
| 52 | 1209 | 12 | 0.01 | EpiAlignment | Astrocytes* |
| 53 | 1209 | 5 | 0.00 | EpiAlignment | Cones (Blue) |
| 54 | 1209 | 7 | 0.01 | EpiAlignment | CD4 T-cells |
| 55 | 1209 | 10 | 0.01 | EpiAlignment | CD8 T-cells |
| 56 | 1209 | 54 | 0.04 | EpiAlignment | Excitatory Camk2a |
| 57 | 1209 | 4 | 0.00 | EpiAlignment | Inhibitory PV |
| 58 | 1209 | 6 | 0.00 | EpiAlignment | Inhibitory VIP |
| 59 | 1209 | 100 | 0.08 | EpiAlignment | Excitatory Layers II-III |
| 60 | 1209 | 58 | 0.05 | EpiAlignment | Excitatory Layer VI* |
| 61 | 1209 | 89 | 0.07 | EpiAlignment | Excitatory DG* |
| 62 | 1209 | 76 | 0.06 | EpiAlignment | Excitatory Layers II-V* |
| 63 | 1209 | 7 | 0.01 | EpiAlignment | Embryonic DA forebrain |
| 64 | 1209 | 92 | 0.08 | EpiAlignment | Inhibitory Gad2 |
| 65 | 1209 | 5 | 0.00 | EpiAlignment | Cones (Green) |
| 66 | 1209 | 38 | 0.03 | EpiAlignment | Inhibitory* |
| 67 | 1209 | 42 | 0.03 | EpiAlignment | Inhibitory MSN* |
| 68 | 1209 | 11 | 0.01 | EpiAlignment | Embryonic DA midbrain |
| 69 | 1209 | 5 | 0.00 | EpiAlignment | Microglia* |
| 70 | 1209 | 9 | 0.01 | EpiAlignment | Neun negative |
| 71 | 1209 | 114 | 0.09 | EpiAlignment | Excitatory Layer VI |
| 72 | 1209 | 4 | 0.00 | EpiAlignment | Oligodendrocytes* |
| 73 | 1209 | 104 | 0.09 | EpiAlignment | Excitatory Layer V |
| 74 | 1209 | 3 | 0.00 | EpiAlignment | Rods |
| 75 | 1209 | 100 | 0.08 | EpiAlignment | Excitatory Layer IV |
| 76 | 715 | 9 | 0.01 | UCSC liftOver | Microglia |
| 77 | 715 | 10 | 0.01 | UCSC liftOver | Astrocytes* |
| 78 | 715 | 9 | 0.01 | UCSC liftOver | Cones (Blue) |
| 79 | 715 | 16 | 0.02 | UCSC liftOver | CD4 T-cells |
| 80 | 715 | 16 | 0.02 | UCSC liftOver | CD8 T-cells |
| 81 | 715 | 71 | 0.10 | UCSC liftOver | Excitatory Camk2a |
| 82 | 715 | 12 | 0.02 | UCSC liftOver | Inhibitory PV |
| 83 | 715 | 13 | 0.02 | UCSC liftOver | Inhibitory VIP |
| 84 | 715 | 109 | 0.15 | UCSC liftOver | Excitatory Layers II-III |
| 85 | 715 | 78 | 0.11 | UCSC liftOver | Excitatory Layer VI* |
| 86 | 715 | 114 | 0.16 | UCSC liftOver | Excitatory DG* |
| 87 | 715 | 114 | 0.16 | UCSC liftOver | Excitatory Layers II-V* |
| 88 | 715 | 17 | 0.02 | UCSC liftOver | Embryonic DA forebrain |
| 89 | 715 | 102 | 0.14 | UCSC liftOver | Inhibitory Gad2 |
| 90 | 715 | 7 | 0.01 | UCSC liftOver | Cones (Green) |
| 91 | 715 | 59 | 0.08 | UCSC liftOver | Inhibitory* |
| 92 | 715 | 66 | 0.09 | UCSC liftOver | Inhibitory MSN* |
| 93 | 715 | 15 | 0.02 | UCSC liftOver | Embryonic DA midbrain |
| 94 | 715 | 6 | 0.01 | UCSC liftOver | Microglia* |
| 95 | 715 | 12 | 0.02 | UCSC liftOver | Neun negative |
| 96 | 715 | 112 | 0.16 | UCSC liftOver | Excitatory Layer VI |
| 97 | 715 | 7 | 0.01 | UCSC liftOver | Oligodendrocytes* |

Continued on next page

**Supplementary Table 6 – continued from previous page**

| | # map | # overlap | precision | method | sample |
|---|---|---|---|---|---|
| 98 | 715 | 109 | 0.15 | UCSC liftOver | Excitatory Layer V |
| 99 | 715 | 5 | 0.01 | UCSC liftOver | Rods |
| 100 | 715 | 99 | 0.14 | UCSC liftOver | Excitatory Layer IV |

Supplementary Table 6: Benchmarking AdaLiftOver on 1,648 SCZ GWAS SNPs.

| | # mapped | # overlapped | precision | method | cell type | trait |
|---|---|---|---|---|---|---|
| 1 | 226 | 14 | 0.06 | AdaLiftOver | B | MCV |
| 2 | 226 | 12 | 0.05 | AdaLiftOver | CD4 | MCV |
| 3 | 226 | 10 | 0.04 | AdaLiftOver | CD8 | MCV |
| 4 | 226 | 39 | 0.17 | AdaLiftOver | CMP | MCV |
| 5 | 226 | 36 | 0.16 | AdaLiftOver | ERY | MCV |
| 6 | 226 | 33 | 0.15 | AdaLiftOver | GMP | MCV |
| 7 | 226 | 16 | 0.07 | AdaLiftOver | Mega | MCV |
| 8 | 226 | 35 | 0.15 | AdaLiftOver | MEP | MCV |
| 9 | 226 | 11 | 0.05 | AdaLiftOver | Mono | MCV |
| 10 | 226 | 14 | 0.06 | AdaLiftOver | NK | MCV |
| 11 | 217 | 12 | 0.06 | bnMapper | B | MCV |
| 12 | 217 | 9 | 0.04 | bnMapper | CD4 | MCV |
| 13 | 217 | 5 | 0.02 | bnMapper | CD8 | MCV |
| 14 | 217 | 36 | 0.17 | bnMapper | CMP | MCV |
| 15 | 217 | 28 | 0.13 | bnMapper | ERY | MCV |
| 16 | 217 | 27 | 0.12 | bnMapper | GMP | MCV |
| 17 | 217 | 18 | 0.08 | bnMapper | Mega | MCV |
| 18 | 217 | 36 | 0.17 | bnMapper | MEP | MCV |
| 19 | 217 | 7 | 0.03 | bnMapper | Mono | MCV |
| 20 | 217 | 11 | 0.05 | bnMapper | NK | MCV |
| 21 | 527 | 18 | 0.03 | EpiAlignment | B | MCV |
| 22 | 527 | 12 | 0.02 | EpiAlignment | CD4 | MCV |
| 23 | 527 | 10 | 0.02 | EpiAlignment | CD8 | MCV |
| 24 | 527 | 31 | 0.06 | EpiAlignment | CMP | MCV |
| 25 | 527 | 31 | 0.06 | EpiAlignment | ERY | MCV |
| 26 | 527 | 29 | 0.06 | EpiAlignment | GMP | MCV |
| 27 | 527 | 15 | 0.03 | EpiAlignment | Mega | MCV |
| 28 | 527 | 31 | 0.06 | EpiAlignment | MEP | MCV |
| 29 | 527 | 12 | 0.02 | EpiAlignment | Mono | MCV |
| 30 | 527 | 16 | 0.03 | EpiAlignment | NK | MCV |
| 31 | 228 | 11 | 0.05 | UCSC liftOver | B | MCV |
| 32 | 228 | 10 | 0.04 | UCSC liftOver | CD4 | MCV |
| 33 | 228 | 6 | 0.03 | UCSC liftOver | CD8 | MCV |
| 34 | 228 | 39 | 0.17 | UCSC liftOver | CMP | MCV |
| 35 | 228 | 30 | 0.13 | UCSC liftOver | ERY | MCV |
| 36 | 228 | 29 | 0.13 | UCSC liftOver | GMP | MCV |
| 37 | 228 | 19 | 0.08 | UCSC liftOver | Mega | MCV |
| 38 | 228 | 38 | 0.17 | UCSC liftOver | MEP | MCV |
| 39 | 228 | 9 | 0.04 | UCSC liftOver | Mono | MCV |
| 40 | 228 | 11 | 0.05 | UCSC liftOver | NK | MCV |
| 41 | 271 | 19 | 0.07 | AdaLiftOver | B | MPV |
| 42 | 271 | 21 | 0.08 | AdaLiftOver | CD4 | MPV |
| 43 | 271 | 22 | 0.08 | AdaLiftOver | CD8 | MPV |
| 44 | 271 | 53 | 0.20 | AdaLiftOver | CMP | MPV |
| 45 | 271 | 29 | 0.11 | AdaLiftOver | ERY | MPV |
| 46 | 271 | 52 | 0.19 | AdaLiftOver | GMP | MPV |
| 47 | 271 | 46 | 0.17 | AdaLiftOver | Mega | MPV |
| 48 | 271 | 39 | 0.14 | AdaLiftOver | MEP | MPV |
| 49 | 271 | 16 | 0.06 | AdaLiftOver | Mono | MPV |

Continued on next page

**Supplementary Table 7 – continued from previous page**

| | # mapped | # overlapped | precision | method | cell type | trait |
|---|---|---|---|---|---|---|
| 50 | 271 | 25 | 0.09 | AdaLiftOver | NK | MPV |
| 51 | 277 | 15 | 0.05 | bnMapper | B | MPV |
| 52 | 277 | 16 | 0.06 | bnMapper | CD4 | MPV |
| 53 | 277 | 19 | 0.07 | bnMapper | CD8 | MPV |
| 54 | 277 | 46 | 0.17 | bnMapper | CMP | MPV |
| 55 | 277 | 20 | 0.07 | bnMapper | ERY | MPV |
| 56 | 277 | 41 | 0.15 | bnMapper | GMP | MPV |
| 57 | 277 | 36 | 0.13 | bnMapper | Mega | MPV |
| 58 | 277 | 34 | 0.12 | bnMapper | MEP | MPV |
| 59 | 277 | 15 | 0.05 | bnMapper | Mono | MPV |
| 60 | 277 | 20 | 0.07 | bnMapper | NK | MPV |
| 61 | 590 | 12 | 0.02 | EpiAlignment | B | MPV |
| 62 | 590 | 8 | 0.01 | EpiAlignment | CD4 | MPV |
| 63 | 590 | 9 | 0.02 | EpiAlignment | CD8 | MPV |
| 64 | 590 | 41 | 0.07 | EpiAlignment | CMP | MPV |
| 65 | 590 | 26 | 0.04 | EpiAlignment | ERY | MPV |
| 66 | 590 | 47 | 0.08 | EpiAlignment | GMP | MPV |
| 67 | 590 | 38 | 0.06 | EpiAlignment | Mega | MPV |
| 68 | 590 | 29 | 0.05 | EpiAlignment | MEP | MPV |
| 69 | 590 | 20 | 0.03 | EpiAlignment | Mono | MPV |
| 70 | 590 | 15 | 0.03 | EpiAlignment | NK | MPV |
| 71 | 287 | 16 | 0.06 | UCSC liftOver | B | MPV |
| 72 | 287 | 16 | 0.06 | UCSC liftOver | CD4 | MPV |
| 73 | 287 | 19 | 0.07 | UCSC liftOver | CD8 | MPV |
| 74 | 287 | 48 | 0.17 | UCSC liftOver | CMP | MPV |
| 75 | 287 | 22 | 0.08 | UCSC liftOver | ERY | MPV |
| 76 | 287 | 42 | 0.15 | UCSC liftOver | GMP | MPV |
| 77 | 287 | 38 | 0.13 | UCSC liftOver | Mega | MPV |
| 78 | 287 | 36 | 0.13 | UCSC liftOver | MEP | MPV |
| 79 | 287 | 16 | 0.06 | UCSC liftOver | Mono | MPV |
| 80 | 287 | 21 | 0.07 | UCSC liftOver | NK | MPV |
| 81 | 161 | 14 | 0.09 | AdaLiftOver | B | Mono |
| 82 | 161 | 17 | 0.11 | AdaLiftOver | CD4 | Mono |
| 83 | 161 | 15 | 0.09 | AdaLiftOver | CD8 | Mono |
| 84 | 161 | 24 | 0.15 | AdaLiftOver | CMP | Mono |
| 85 | 161 | 8 | 0.05 | AdaLiftOver | ERY | Mono |
| 86 | 161 | 39 | 0.24 | AdaLiftOver | GMP | Mono |
| 87 | 161 | 23 | 0.14 | AdaLiftOver | Mega | Mono |
| 88 | 161 | 10 | 0.06 | AdaLiftOver | MEP | Mono |
| 89 | 161 | 29 | 0.18 | AdaLiftOver | Mono | Mono |
| 90 | 161 | 20 | 0.12 | AdaLiftOver | NK | Mono |
| 91 | 186 | 14 | 0.08 | bnMapper | B | Mono |
| 92 | 186 | 10 | 0.05 | bnMapper | CD4 | Mono |
| 93 | 186 | 9 | 0.05 | bnMapper | CD8 | Mono |
| 94 | 186 | 22 | 0.12 | bnMapper | CMP | Mono |
| 95 | 186 | 7 | 0.04 | bnMapper | ERY | Mono |
| 96 | 186 | 41 | 0.22 | bnMapper | GMP | Mono |
| 97 | 186 | 29 | 0.16 | bnMapper | Mega | Mono |

Continued on next page

**Supplementary Table 7 – continued from previous page**

| | # mapped | # overlapped | precision | method | cell type | trait |
|---|---|---|---|---|---|---|
| 98 | 186 | 11 | 0.06 | bnMapper | MEP | Mono |
| 99 | 186 | 24 | 0.13 | bnMapper | Mono | Mono |
| 100 | 186 | 16 | 0.09 | bnMapper | NK | Mono |
| 101 | 383 | 14 | 0.04 | EpiAlignment | B | Mono |
| 102 | 383 | 9 | 0.02 | EpiAlignment | CD4 | Mono |
| 103 | 383 | 15 | 0.04 | EpiAlignment | CD8 | Mono |
| 104 | 383 | 12 | 0.03 | EpiAlignment | CMP | Mono |
| 105 | 383 | 3 | 0.01 | EpiAlignment | ERY | Mono |
| 106 | 383 | 26 | 0.07 | EpiAlignment | GMP | Mono |
| 107 | 383 | 10 | 0.03 | EpiAlignment | Mega | Mono |
| 108 | 383 | 9 | 0.02 | EpiAlignment | MEP | Mono |
| 109 | 383 | 20 | 0.05 | EpiAlignment | Mono | Mono |
| 110 | 383 | 10 | 0.03 | EpiAlignment | NK | Mono |
| 111 | 202 | 16 | 0.08 | UCSC liftOver | B | Mono |
| 112 | 202 | 11 | 0.05 | UCSC liftOver | CD4 | Mono |
| 113 | 202 | 9 | 0.04 | UCSC liftOver | CD8 | Mono |
| 114 | 202 | 24 | 0.12 | UCSC liftOver | CMP | Mono |
| 115 | 202 | 7 | 0.03 | UCSC liftOver | ERY | Mono |
| 116 | 202 | 44 | 0.22 | UCSC liftOver | GMP | Mono |
| 117 | 202 | 31 | 0.15 | UCSC liftOver | Mega | Mono |
| 118 | 202 | 11 | 0.05 | UCSC liftOver | MEP | Mono |
| 119 | 202 | 25 | 0.12 | UCSC liftOver | Mono | Mono |
| 120 | 202 | 17 | 0.08 | UCSC liftOver | NK | Mono |
| 121 | 148 | 31 | 0.21 | AdaLiftOver | B | Lymph |
| 122 | 148 | 31 | 0.21 | AdaLiftOver | CD4 | Lymph |
| 123 | 148 | 33 | 0.22 | AdaLiftOver | CD8 | Lymph |
| 124 | 148 | 16 | 0.11 | AdaLiftOver | CMP | Lymph |
| 125 | 148 | 12 | 0.08 | AdaLiftOver | ERY | Lymph |
| 126 | 148 | 16 | 0.11 | AdaLiftOver | GMP | Lymph |
| 127 | 148 | 18 | 0.12 | AdaLiftOver | Mega | Lymph |
| 128 | 148 | 15 | 0.10 | AdaLiftOver | MEP | Lymph |
| 129 | 148 | 13 | 0.09 | AdaLiftOver | Mono | Lymph |
| 130 | 148 | 27 | 0.18 | AdaLiftOver | NK | Lymph |
| 131 | 194 | 20 | 0.10 | bnMapper | B | Lymph |
| 132 | 194 | 23 | 0.12 | bnMapper | CD4 | Lymph |
| 133 | 194 | 24 | 0.12 | bnMapper | CD8 | Lymph |
| 134 | 194 | 9 | 0.05 | bnMapper | CMP | Lymph |
| 135 | 194 | 7 | 0.04 | bnMapper | ERY | Lymph |
| 136 | 194 | 8 | 0.04 | bnMapper | GMP | Lymph |
| 137 | 194 | 10 | 0.05 | bnMapper | Mega | Lymph |
| 138 | 194 | 11 | 0.06 | bnMapper | MEP | Lymph |
| 139 | 194 | 8 | 0.04 | bnMapper | Mono | Lymph |
| 140 | 194 | 18 | 0.09 | bnMapper | NK | Lymph |
| 141 | 412 | 19 | 0.05 | EpiAlignment | B | Lymph |
| 142 | 412 | 20 | 0.05 | EpiAlignment | CD4 | Lymph |
| 143 | 412 | 19 | 0.05 | EpiAlignment | CD8 | Lymph |
| 144 | 412 | 13 | 0.03 | EpiAlignment | CMP | Lymph |
| 145 | 412 | 5 | 0.01 | EpiAlignment | ERY | Lymph |

Continued on next page

**Supplementary Table 7 – continued from previous page**

| | # mapped | # overlapped | precision | method | cell type | trait |
|---|---|---|---|---|---|---|
| 146 | 412 | 12 | 0.03 | EpiAlignment | GMP | Lymph |
| 147 | 412 | 11 | 0.03 | EpiAlignment | Mega | Lymph |
| 148 | 412 | 4 | 0.01 | EpiAlignment | MEP | Lymph |
| 149 | 412 | 8 | 0.02 | EpiAlignment | Mono | Lymph |
| 150 | 412 | 20 | 0.05 | EpiAlignment | NK | Lymph |
| 151 | 209 | 22 | 0.11 | UCSC liftOver | B | Lymph |
| 152 | 209 | 25 | 0.12 | UCSC liftOver | CD4 | Lymph |
| 153 | 209 | 26 | 0.12 | UCSC liftOver | CD8 | Lymph |
| 154 | 209 | 11 | 0.05 | UCSC liftOver | CMP | Lymph |
| 155 | 209 | 7 | 0.03 | UCSC liftOver | ERY | Lymph |
| 156 | 209 | 9 | 0.04 | UCSC liftOver | GMP | Lymph |
| 157 | 209 | 11 | 0.05 | UCSC liftOver | Mega | Lymph |
| 158 | 209 | 11 | 0.05 | UCSC liftOver | MEP | Lymph |
| 159 | 209 | 10 | 0.05 | UCSC liftOver | Mono | Lymph |
| 160 | 209 | 20 | 0.10 | UCSC liftOver | NK | Lymph |
| 161 | 47 | 1 | 0.02 | AdaLiftOver | B | Alzheimer |
| 162 | 47 | 0 | 0.00 | AdaLiftOver | CD4 | Alzheimer |
| 163 | 47 | 3 | 0.06 | AdaLiftOver | CD8 | Alzheimer |
| 164 | 47 | 3 | 0.06 | AdaLiftOver | CMP | Alzheimer |
| 165 | 47 | 1 | 0.02 | AdaLiftOver | ERY | Alzheimer |
| 166 | 47 | 3 | 0.06 | AdaLiftOver | GMP | Alzheimer |
| 167 | 47 | 3 | 0.06 | AdaLiftOver | Mega | Alzheimer |
| 168 | 47 | 1 | 0.02 | AdaLiftOver | MEP | Alzheimer |
| 169 | 47 | 0 | 0.00 | AdaLiftOver | Mono | Alzheimer |
| 170 | 47 | 1 | 0.02 | AdaLiftOver | NK | Alzheimer |
| 171 | 91 | 0 | 0.00 | bnMapper | B | Alzheimer |
| 172 | 91 | 0 | 0.00 | bnMapper | CD4 | Alzheimer |
| 173 | 91 | 2 | 0.02 | bnMapper | CD8 | Alzheimer |
| 174 | 91 | 2 | 0.02 | bnMapper | CMP | Alzheimer |
| 175 | 91 | 0 | 0.00 | bnMapper | ERY | Alzheimer |
| 176 | 91 | 3 | 0.03 | bnMapper | GMP | Alzheimer |
| 177 | 91 | 4 | 0.04 | bnMapper | Mega | Alzheimer |
| 178 | 91 | 0 | 0.00 | bnMapper | MEP | Alzheimer |
| 179 | 91 | 0 | 0.00 | bnMapper | Mono | Alzheimer |
| 180 | 91 | 0 | 0.00 | bnMapper | NK | Alzheimer |
| 181 | 204 | 6 | 0.03 | EpiAlignment | B | Alzheimer |
| 182 | 204 | 3 | 0.01 | EpiAlignment | CD4 | Alzheimer |
| 183 | 204 | 0 | 0.00 | EpiAlignment | CD8 | Alzheimer |
| 184 | 204 | 5 | 0.02 | EpiAlignment | CMP | Alzheimer |
| 185 | 204 | 2 | 0.01 | EpiAlignment | ERY | Alzheimer |
| 186 | 204 | 6 | 0.03 | EpiAlignment | GMP | Alzheimer |
| 187 | 204 | 2 | 0.01 | EpiAlignment | Mega | Alzheimer |
| 188 | 204 | 2 | 0.01 | EpiAlignment | MEP | Alzheimer |
| 189 | 204 | 1 | 0.00 | EpiAlignment | Mono | Alzheimer |
| 190 | 204 | 3 | 0.01 | EpiAlignment | NK | Alzheimer |
| 191 | 94 | 0 | 0.00 | UCSC liftOver | B | Alzheimer |
| 192 | 94 | 0 | 0.00 | UCSC liftOver | CD4 | Alzheimer |
| 193 | 94 | 2 | 0.02 | UCSC liftOver | CD8 | Alzheimer |

Continued on next page

**Supplementary Table 7 – continued from previous page**

| | # mapped | # overlapped | precision | method | cell type | trait |
|---|---|---|---|---|---|---|
| 194 | 94 | 2 | 0.02 | UCSC liftOver | CMP | Alzheimer |
| 195 | 94 | 0 | 0.00 | UCSC liftOver | ERY | Alzheimer |
| 196 | 94 | 3 | 0.03 | UCSC liftOver | GMP | Alzheimer |
| 197 | 94 | 4 | 0.04 | UCSC liftOver | Mega | Alzheimer |
| 198 | 94 | 0 | 0.00 | UCSC liftOver | MEP | Alzheimer |
| 199 | 94 | 0 | 0.00 | UCSC liftOver | Mono | Alzheimer |
| 200 | 94 | 0 | 0.00 | UCSC liftOver | NK | Alzheimer |

Supplementary Table 7: Benchmarking AdaLiftOver on hematopoietic GWAS SNPs for 4 traits: MCV (570), MPV (646), Mono (441), Lymph (466). The Alzheimer (218) GWAS SNPs are treated as negative controls.

|  | # mapped GWAS | # overlapped GWAS | # genes | precision | method |
|---|---|---|---|---|---|
| 1 | 1060 | 86 | 30 | 0.08 | AdaLiftOver |
| 2 | 1502 | 62 | 28 | 0.04 | bnMapper |
| 3 | 1586 | 65 | 29 | 0.04 | UCSC liftOver |
| 4 | 2526 | 62 | 23 | 0.02 | EpiAlignment |

Supplementary Table 8: Benchmarking AdaLiftOver on BMD GWAS SNPs. Mapping of the 3,125 BMD fine-mapped GWAS SNPs to 52 BMD genes in mouse. Distance = 250 kb.

|  | # mapped GWAS | # overlapped GWAS | # genes | precision | distance | method |
|---|---|---|---|---|---|---|
| 1 | 29486 | 14 | 11 | 0.00 | 0 | AdaLiftOver |
| 2 | 38797 | 5 | 3 | 0.00 | 0 | bnMapper |
| 3 | 41278 | 6 | 4 | 0.00 | 0 | UCSC liftOver |
| 4 | 29486 | 1096 | 71 | 0.04 | 100kb | AdaLiftOver |
| 5 | 38797 | 791 | 67 | 0.02 | 100kb | bnMapper |
| 6 | 41278 | 878 | 74 | 0.02 | 100kb | UCSC liftOver |
| 7 | 29486 | 2140 | 90 | 0.07 | 250kb | AdaLiftOver |
| 8 | 38797 | 1661 | 89 | 0.04 | 250kb | bnMapper |
| 9 | 41278 | 1813 | 102 | 0.04 | 250kb | UCSC liftOver |

Supplementary Table 9: Benchmarking AdaLiftOver on BMD GWAS SNPs. Mapping of the 116,402 BMD fine-mapped GWAS SNPs to 200 BMD genes in mouse.

#### 3 Supplementary Figures

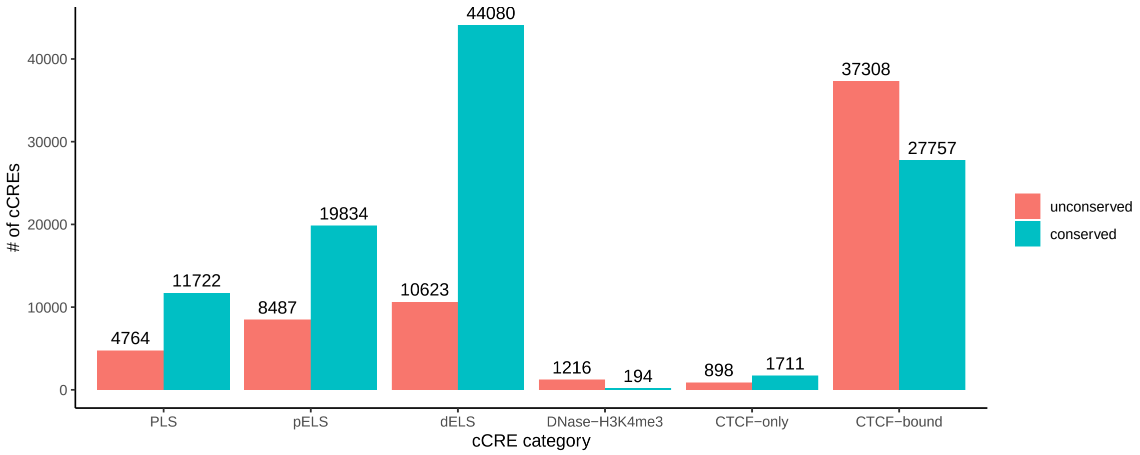

Supplementary Figure 1: The summary of categories between orthologous cCREs. X-axis: the human cCRE category. Y-axis: the number of orthologous mouse cCREs that fall into the same category (conserved) or a different category (unconserved).

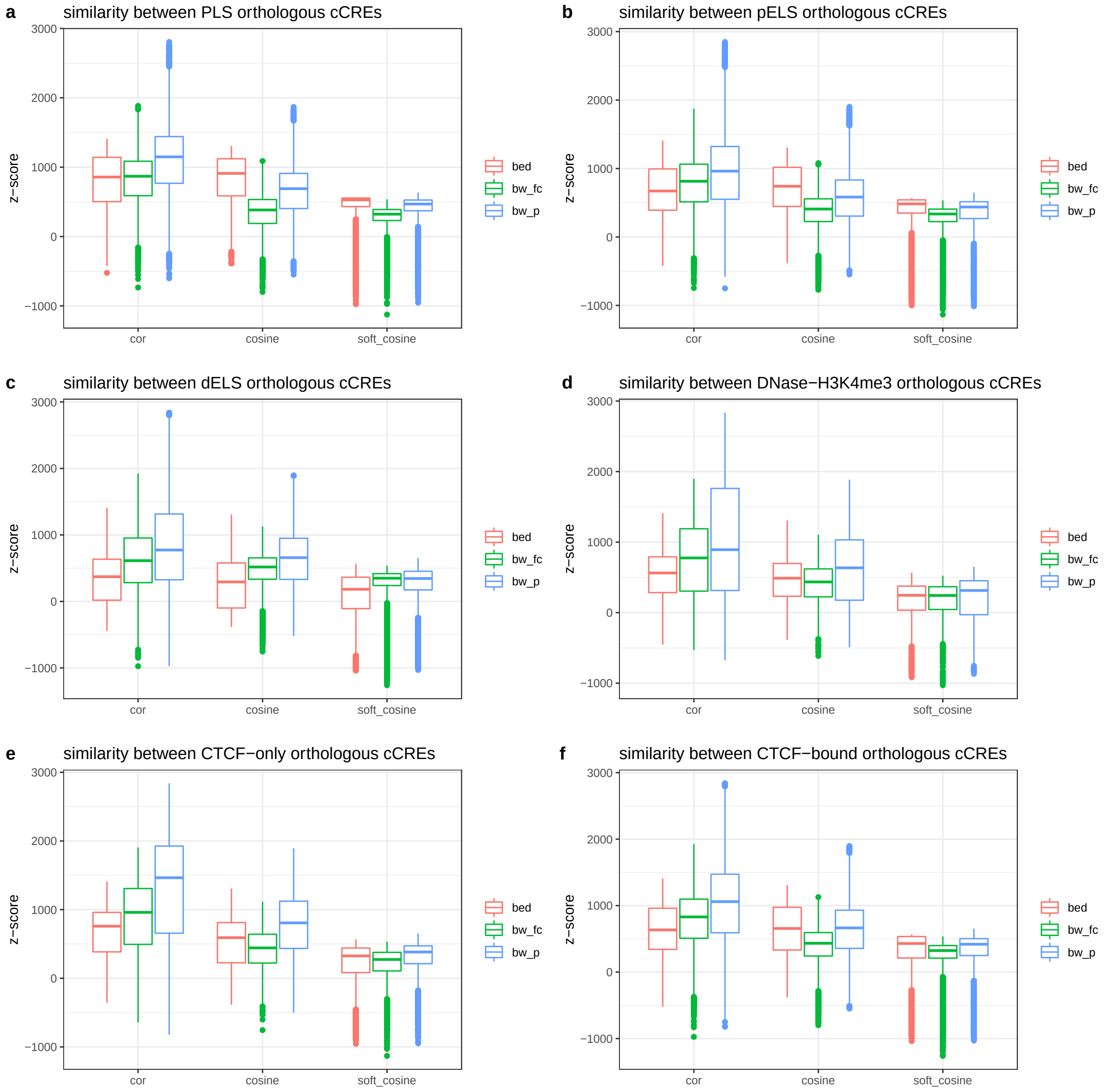

Supplementary Figure 2: Comparison of epigenome similarities for different classes of orthogolous cCREs with three different metrics (cor: Pearson's correlation; cosine: cosine similarity; soft_cosine: soft cosine similarity) and three different data formats (bed: BED format; bw_fc: fold change BigWig format; bw_p: p-value BigWig format) for constructing the epigenomic features. A null distribution for each of the similarity scores is estimated by randomly permuting cCREs 10,000 times. Observed similarity scores were transformed into z-scores using the mean and variance estimates of these null distributions.

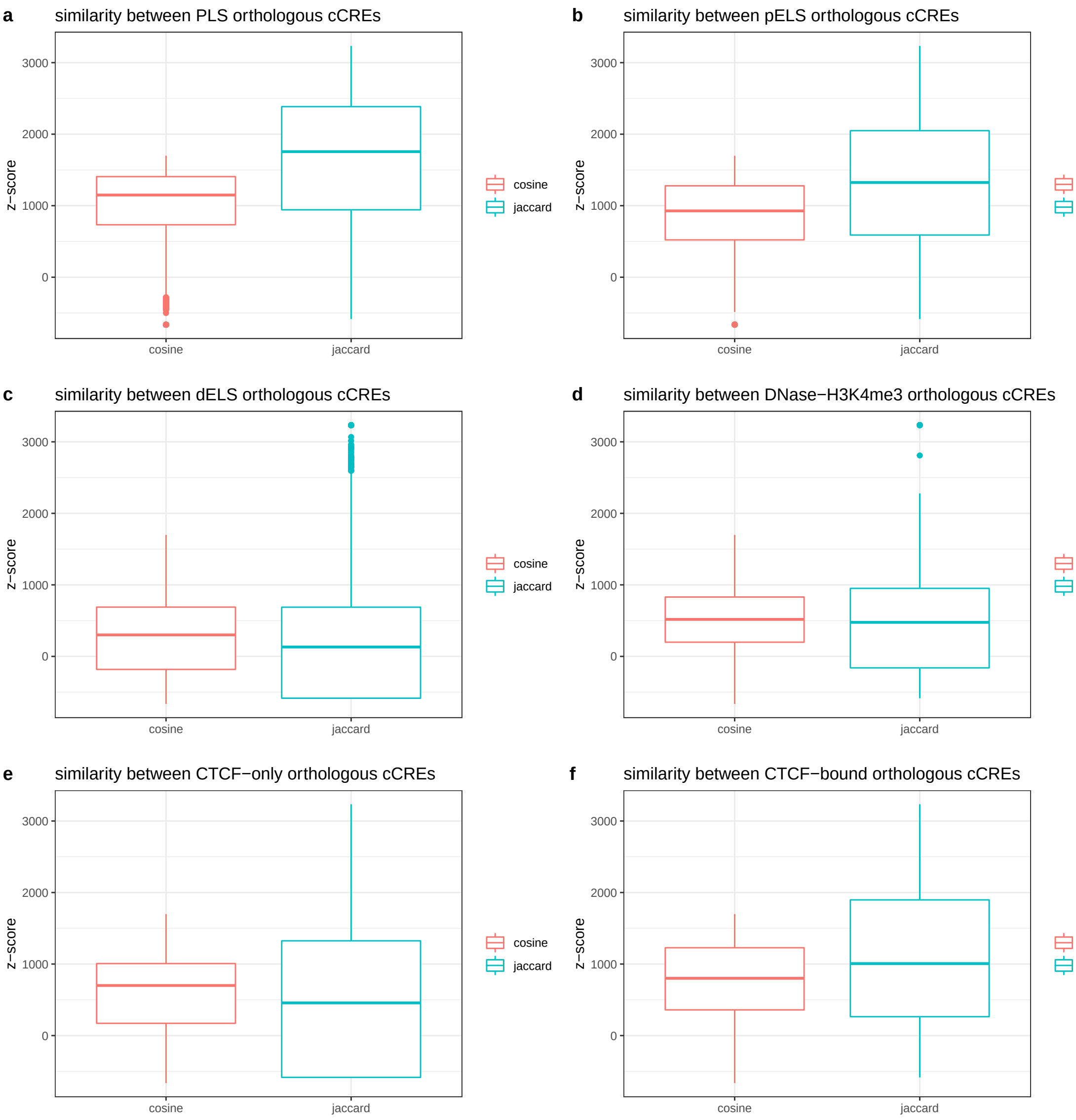

Supplementary Figure 3: Comparison of similarity metrics (cosine: cosine similarity; jaccard: Jaccard similarity) for binary epigenomic features (BED format). A null distribution for each of the similarity scores is estimated by randomly permuting cCREs. Observed similarity scores were transformed into z-scores using the mean and variance estimates of these null distributions.

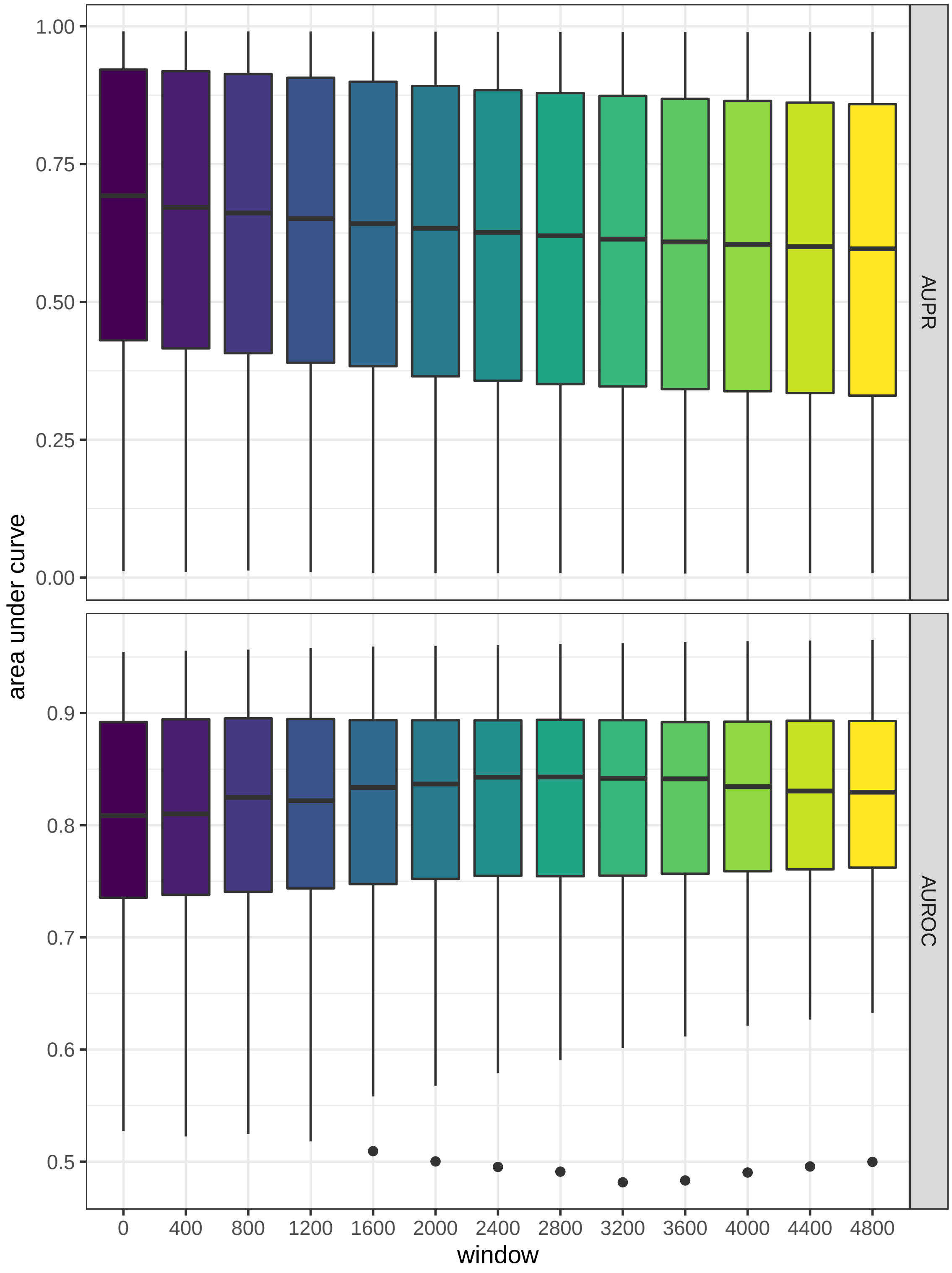

Supplementary Figure 4: Summary of the leave-one-out cross-validation experiments with 67 epigenome datasets over a grid of windows.

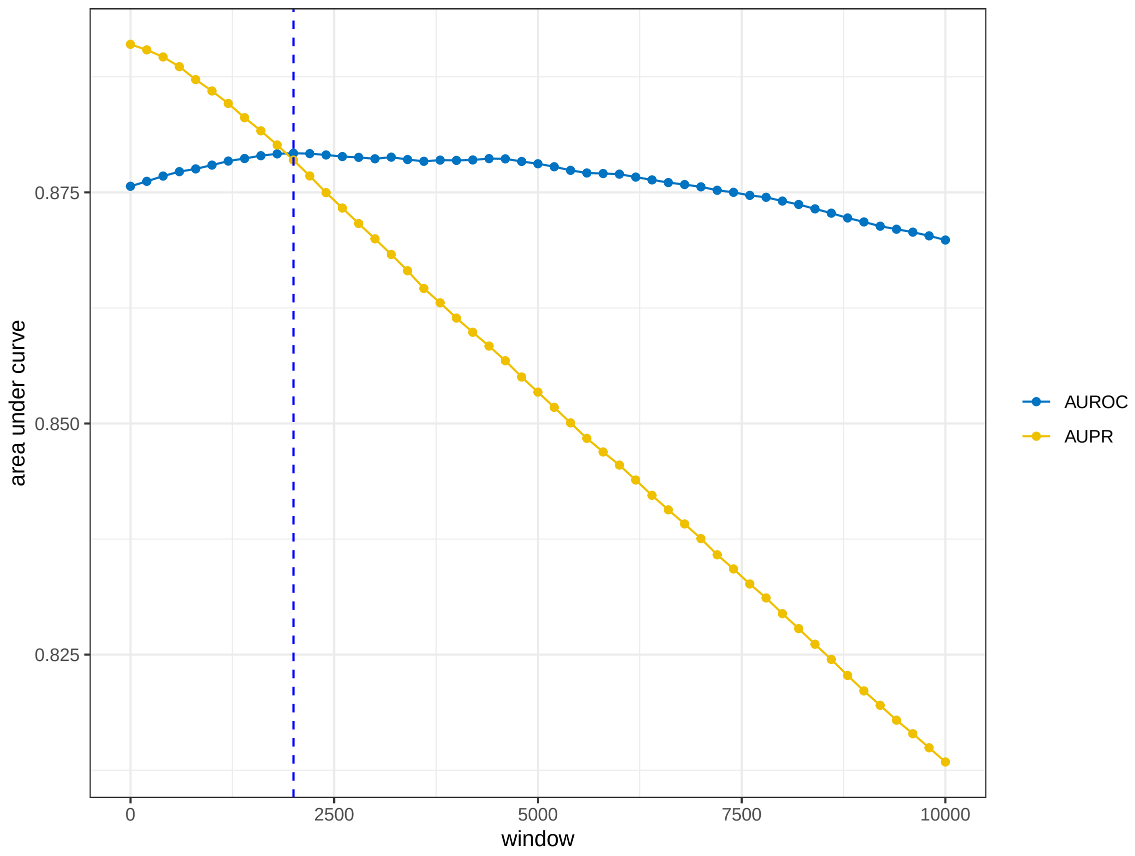

Supplementary Figure 5: The area under ROC/PR curve over a grid of windows from the mouse islet ATAC-seq peak analysis. AUROC and AUPR values are obtained for the set of candidate target regions generated at different local window sizes by fitting a logistic regression on the data labelled using the human islet ATAC-seq peaks as the gold standard. The vertical blue dashed line depicts window size of 2 kb.

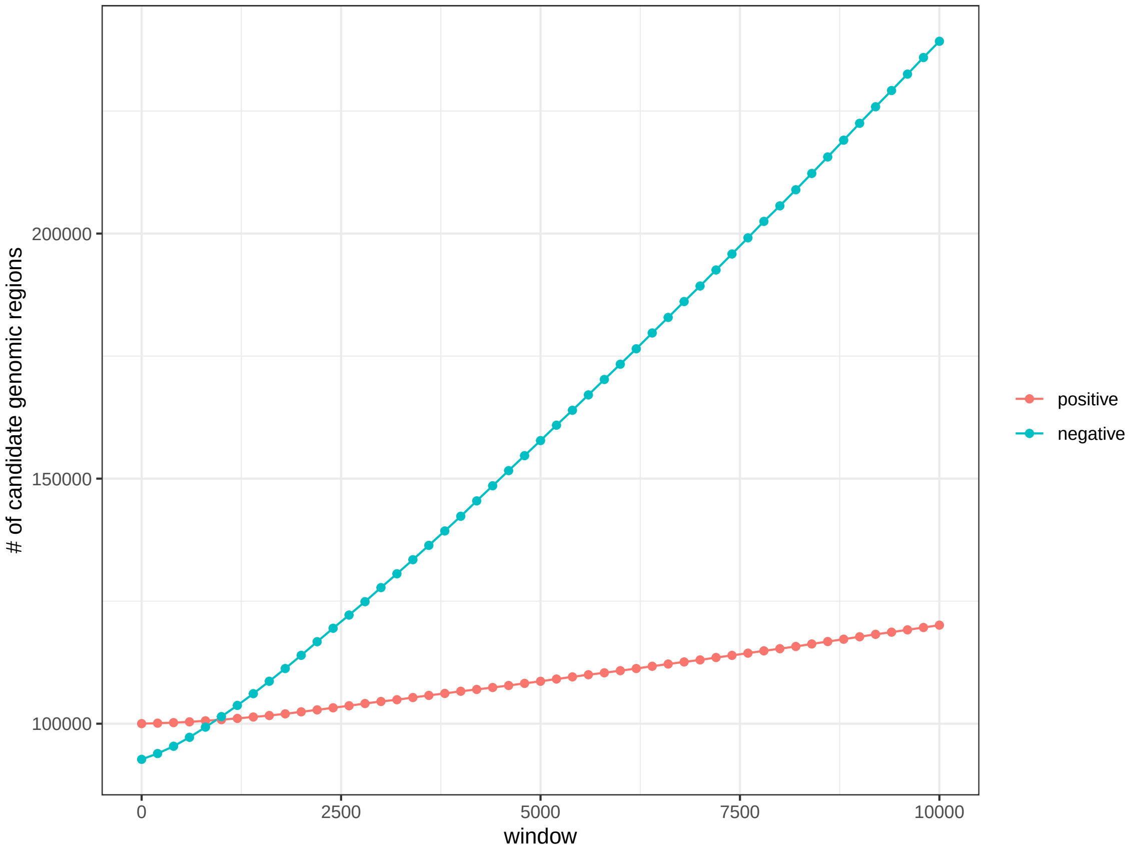

Supplementary Figure 6: As the local window size increases, it generates more candidate target regions, which results in an imbalance between the numbers of positive and negative cases, and hence the decreasing pattern in AUPR (Supplementary Figure 5).

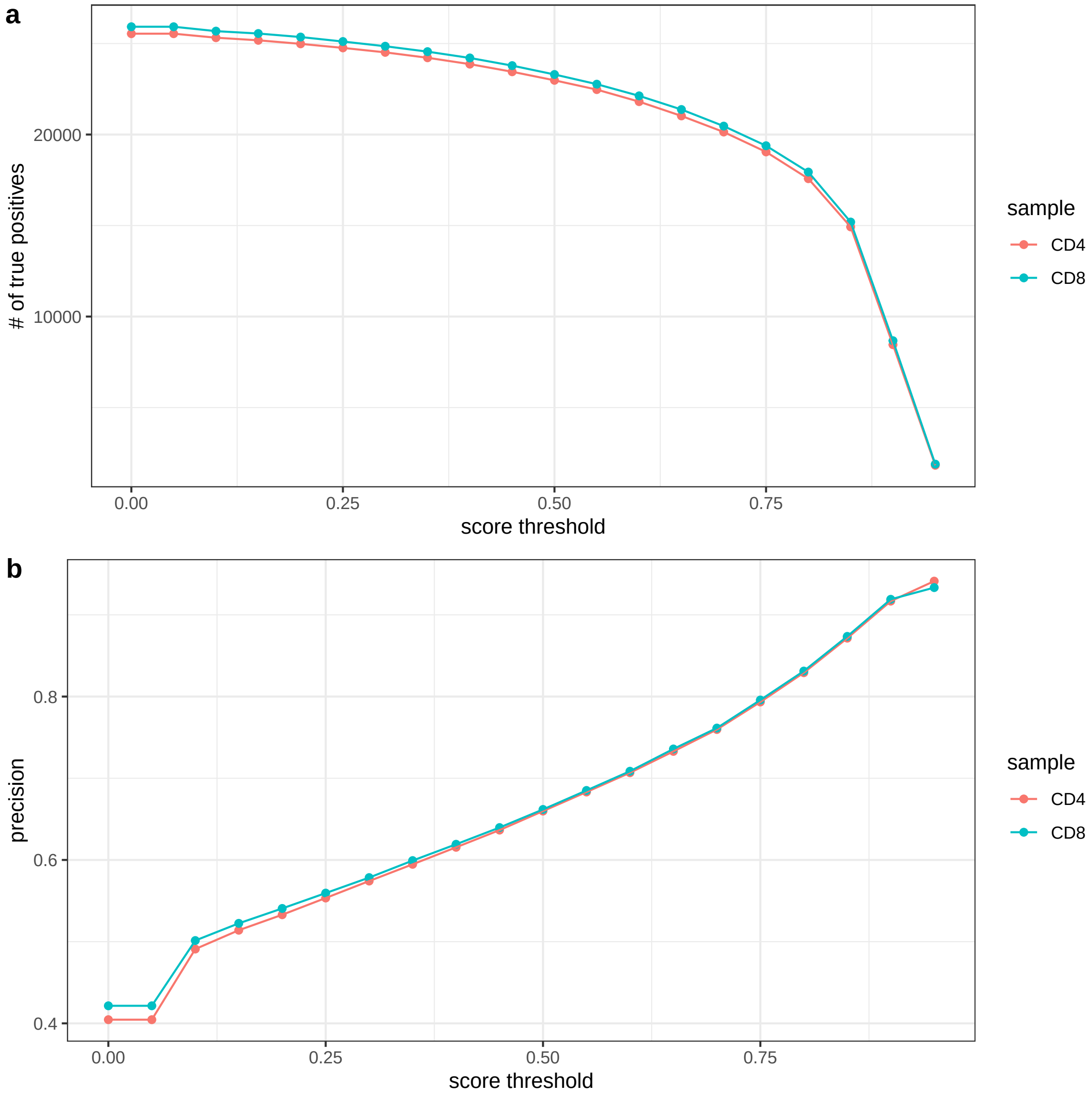

Supplementary Figure 7: The effects of logistic probability score threshold over # of true positives (a) and precision (b) for the orthologous CD4 and CD8 ATAC-seq data. # of true positives: the number of mouse-derived peaks that overlaps human peaks; precision: # of true positives / # of mouse-derived peaks.

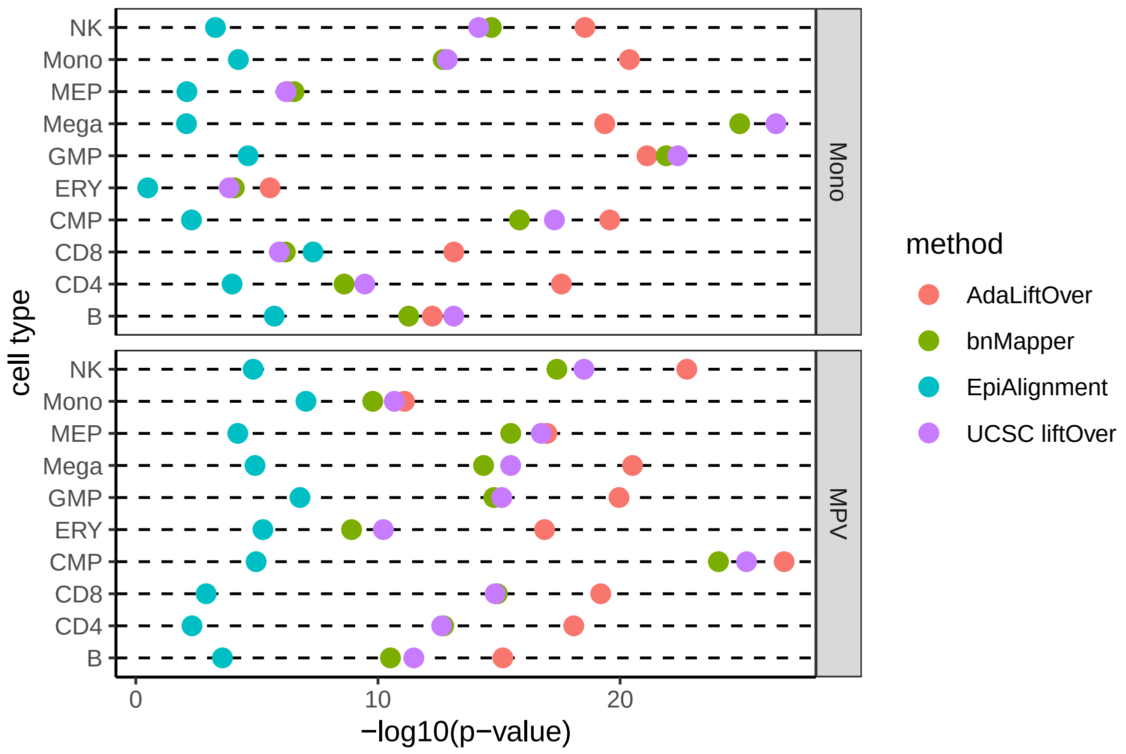

Supplementary Figure 8: Enrichment analysis for fine-mapped GWAS SNPs from two other hematopoietic traits with respect to 10 hematopoietic ATAC-seq peaks. Mono: monocyte count, MPV: mean platelet volume.

#### References

Cheng, Y., Ma, Z., Kim, B.-H., Wu, W., Cayting, P., Boyle, A. P., Sundaram, V., Xing, X., Dogan, N., Li, J., *et al.* (2014). Principles of regulatory information conservation between mouse and human. *Nature*, **515**(7527), 371–375.

Hook, P. W. and McCallion, A. S. (2020). Leveraging mouse chromatin data for heritability enrichment informs common disease architecture and reveals cortical layer contributions to schizophrenia. *Genome research*, **30**(4), 528–539.

Morris, J. A., Kemp, J. P., Youlten, S. E., Laurent, L., Logan, J. G., Chai, R. C., Vulpescu, N. A., Forgetta, V., Kleinman, A., Mohanty, S. T., *et al.* (2019). An atlas of genetic influences on osteoporosis in humans and mice. *Nature genetics*, **51**(2), 258–266.

Ulirsch, J. C., Lareau, C. A., Bao, E. L., Ludwig, L. S., Guo, M. H., Benner, C., Satpathy, A. T., Kartha, V. K., Salem, R. M., Hirschhorn, J. N., *et al.* (2019). Interrogation of human hematopoiesis at single-cell and single-variant resolution. *Nature genetics*, **51**(4), 683–693.

Xiang, G., Keller, C. A., Heuston, E., Giardine, B. M., An, L., Wixom, A. Q., Miller, A., Cockburn, A., Sauria, M. E., Weaver, K., *et al.* (2020). An integrative view of the regulatory and transcriptional landscapes in mouse hematopoiesis. *Genome research*, **30**(3), 472–484.
